# Imaging mass cytometry reveals early β-cell dysfunction and changes in immune signatures during type 1 diabetes progression in human pancreata

**DOI:** 10.1101/2025.03.05.641526

**Authors:** Nathan Steenbuck, Nicolas Damond, Stefanie Engler, Irina Kusmartseva, Amanda L. Posgai, Denise M. Drotar, MacKenzie D. Williams, Natalie de Souza, Todd M. Brusko, Maigan A. Brusko, Clive H. Wasserfall, Mark A. Atkinson, Bernd Bodenmiller

## Abstract

The natural history and pathogenesis of type 1 diabetes, particularly during the autoantibody-positive stages preceding clinical onset, are not well understood, in part, due to limited availability of human pancreatic samples. Here, we studied 88 organ donors, including 28 single autoantibody-positive and 10 multiple autoantibody-positive donors, by imaging mass cytometry. Approximately 10,000 islets and 16 million single-cells were spatially analyzed using 79 antibodies revealing both β-cell states and the islet-immune interface. We identified IAPP loss from β-cells as an indicator of pre-clinical disease. Alterations in Interferon signatures and downregulation across lineage and functional markers, including markers of endoplasmic reticulum stress, were characteristic of recent-onset disease. Further, in single autoantibody-positive donors, we identified pro-inflammatory myeloid cells and PD1^+^ memory CD4^+^ T cells, and in multiple autoantibody-positive samples, found islet-specific and exhausted-like effector CD8^+^ T cells. Multiple immune cell subtypes were associated with young age, disease severity and insulitis. This dataset is a major step toward creation of a multi-modal type 1 diabetes disease atlas that will be useful for identifying potential drug targets and association of disease features with clinical co-variates and trial outcomes.

## Introduction

Type 1 diabetes (T1D) is a chronic autoimmune disease characterized by the selective loss of insulin-producing pancreatic β-cells^1^. Despite the high global disease burden, neither a cure nor a preventive strategy has been identified in part due to an incomplete understanding of T1D pathogenesis^2,3^. β-cell loss is triggered by an unknown autoimmune event that results in development of autoantibodies that target β-cell antigens. These autoantibodies can emerge years prior to clinical symptoms and offer valuable prognostic insights. The presence of a single autoantibody (sAAb+) indicates a lifetime risk of approximately 15% of developing T1D, whereas the presence of two or more autoantibodies (mAAb+) signals a significantly higher lifetime risk of approximately 85%^4^. mAAb+ subjects are classified as having stage 1, and mAAb+ and dysglycemic subjects as having stage 2 disease^5^. Stage 3 T1D is typically diagnosed upon significant loss of β-cell mass, and symptoms at diagnosis and time to complete β-cell loss are highly variable between patients^5,6^. Understanding the biological processes driving T1D progression, particularly during the sAAb+ and mAAb+ stages, will be essential to guide development of therapeutic strategies that preserve endogenous β-cells.

T1D is considered to be mainly a disease of the adaptive immune system^7^ given the importance of autoantibodies, the observed β-cell destruction by autoreactive T cells^8,9^ and the partial success of T cell-targeting therapeutics in delaying T1D onset^10,11,12,13,14,15^. Mouse studies indicate that components of the innate immune system, particularly dendritic cells (DCs) and macrophages, influence initiation^16,17^ and progression of the disease^18,19,20,21^, but the role of the innate immune system in the human pancreas remains poorly defined^22^. T1D is also associated with dysregulation in the endocrine compartment, including MHC-I hyperexpression^23,24^, secretion and response to pro-inflammatory cytokines^25^, and responses to endoplasmic reticulum (ER) stress by β-cells^26,27,28^. Given the wide range of disease states at the pancreatic islet-immune interface and heterogeneity associated with age, clinical presentation, and intervention response^29^, a comprehensive characterization of human pancreata by single-cell, spatially resolved, and multiplexed modalities at large scale and across ages is needed.

So far, only a few spatial multiplexed protein imaging studies have been conducted on human T1D samples, and these studies were limited by the small number of donors overall with just two autoantibody positive donors evaluated to date^6,30,31,32^. To gain insights into the earliest steps of T1D development, we used imaging mass cytometry (IMC) on pancreatic sections from 88 organ donors spanning the entire spectrum of T1D disease progression, including 28 sAAb+ and 10 mAAb+ donors, and controls without T1D. We used two 45-plex antibody panels to extensively characterize the islet-immune interface. Using these IMC data, we identified IAPP loss, pro-inflammatory myeloid phenotypes, and PD1^+^ T cells as critical indicators of progression in mAAb+ donors, suggesting that these are potential therapeutic targets. Further, we integrated our data with clinical co-variates to describe disease phenotypes associated with age.

## Results

### Highly multiplex imaging of human T1D samples

Studies of early T1D progression have been limited by sample availability^3^. For this study, we obtained pancreatic tissue sections from 88 cadaveric organ donors across the entire spectrum of T1D risk and progression from the Network for Pancreatic Organ donors with Diabetes (nPOD) biorepository (Figure 1A, Table S1). Included were samples from sAAb+ (N=28) and mAAb+ (N=10) donors, recent-onset (≤ 2 years) (Onset) T1D donors (N=21), long-duration (≥ 3 years) (LD) T1D donors (N=14) as well as controls (N=15) matched by age, sex, and BMI (Figure S1). Enabling the study of potential age-associated endotypes^29^, 11 donors were less than 7 years of age, and 9 were between 7 and 12 years of age. For a more robust statistical analysis, we pooled donors into groups of <13 years of age (N=20) and ≥13 years of age (N=68).

**Figure 1:**
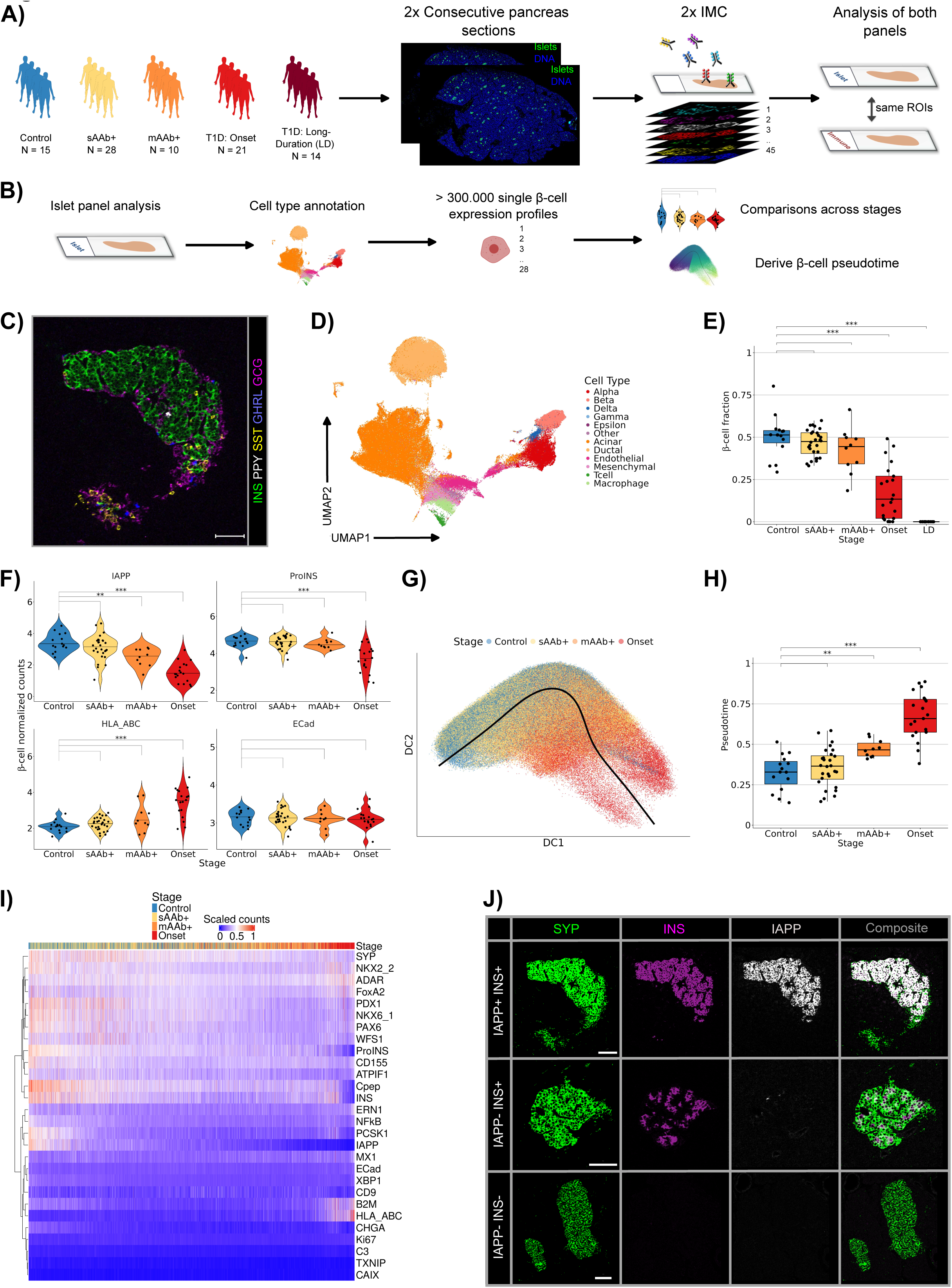
IAPP is downregulated and MHC-I is upregulated along the inferred β-cell pseudotime. **A:** Pancreatic samples from 88 cadaveric organ donors were stained with two 45-plex antibody panels on consecutive sections, and corresponding ROIs (n=75 per sample) were analyzed by imaging mass cytometry. Single cells and islets were segmented, and the resulting data analyzed. **B:** Data analysis of the islet panel. **C:** Representative image of a pancreatic islet from a non-diabetic control donor. INS, GCG, SST, PPY, and GHRL are markers of β-, α-, δ-, γ-, and ε-cells, respectively. **D:** UMAP of a subset of 190,000 cells colored by cell type. **E:** Fraction of β-cells per islet in control and disease stage groups. Each dot indicates the average β-cell fraction per sample. **F:** The expression of the indicated β-cell markers in control and disease stage groups. ECad was included as its expression is expected to be similar in all groups. Each dot indicates data from one donor with expression averaged across ROIs. **G:** Trajectory inferred using *slingshot* from expression profiles of single β-cells projected onto a diffusion map. β-cells are colored by donor stage. **H:** Average pseudotime of β-cells in control and disease stage groups. Each dot indicates the average per sample. **I:** Heatmap of expression of lineage and functional markers expressed in β-cells with columns representing ROIs arranged according to inferred pseudotime progression. Scaled expression values are depicted. **J:** Representative images of samples from control and disease stages stained for SYP (islets), IAPP, and INS. Islets from control donors were IAPP+INS+ and islets from long-duration (LD) T1D donors were IAPP-INS-. IAPP-INS+ were observed in multiple ROIs of mAAb+ and Onset donors. Scale bar is 100 µm. Tests: Linear mixed- effects models with random intercept by organ donor were used to compare differential expression between disease stages. Differential abundance between disease stages was computed using *edgeR*.^∗^ *P* <0.05; ^∗∗^ *P* <0.01; ^∗∗∗^ *P* <0.001. Significance was determined for each stage by testing for differential abundance against controls using *edgeR* and by testing for differential expression against controls by fitting a linear mixed-effects model.

To extensively analyze the endocrine and auto-immune compartments as well as the islet-immune interface, we designed two IMC antibody panels targeting a total of 79 protein markers (Table S2, S3). We selected these antibodies by testing their specificity and association to T1D progression in a pilot study of 130 antibodies applied to Control (N=4), sAAb+ (N=4), mAAb+ (N=4), and Onset T1D donors (N=4) (Table S4-8). Using these two optimized panels, we imaged two 4-µm-thick consecutive sections from each donor. Because the speed of IMC data acquisition limits imaging of whole-slide pancreas sections, we used immunofluorescence imaging with antibodies against islet markers (CD99) and immune markers (Islet panel: CD45RO/CD45RA; Immune panel: CD3e) to pre-select regions of interest (ROIs). We chose ROIs to comprise islets with proximal exocrine tissue, with the goal of capturing both the endocrine state and the immune compartment. We then stained the sections with metal-labeled primary antibodies and acquired 75 selected ROIs per section by IMC. The same locations were imaged in consecutive sections to facilitate an integrative analysis by both panels. We segmented cells and islets in each image (Figure S2). Islet segmentation was performed using a custom UNet-based convolutional neural network. Using the islet masks, we extracted distances to islet-edge measurements for each cell to study islet infiltration by immune cells. After stringent quality control and channel signal spillover correction (Figure S2D), our dataset comprised 7,025 ROIs for each antibody panel, 10,413 islets, and >16 million cells. To our knowledge, this represents, by far, the largest multiplexed imaging effort of T1D samples to date.

### β-cells and functional β-cell markers are lost during disease progression

One of the antibody panels was designed to study dysregulation in the endocrine compartment and included antibodies to lineage markers of islet cells as well as functional markers of ER stress (WFS-1, IRE1a, XBP1), the Interferon (IFN) response (MX1, ADAR), islet inflammation (B2M, HLA-ABC), and redox and inflammatory signaling (TXNIP, NF-κB, CD155) (Table S3). We used this panel to annotate 11 cell types, to compare single-cell β-cell protein expression levels across 28 included β-cell markers, and to infer pseudotime ordering of β-cells (Figure 1B). We annotated cells by clustering single-cell lineage marker expression profiles and then, implementing a multi-step annotation approach for the resulting clusters. First, we annotated abundant (α, β, δ) and rare (ε, γ) endocrine cell types (Figure 1C) and subsequently labeled all non-endocrine cells as acinar, ductal, endothelial, mesenchymal, macrophage, T cell, or ‘other’ (Figure 1D, Figure S3A). The last category likely encompasses immune cells such as neutrophils. In total, we annotated over 6.2 million cells with the islet panel. Of the 975,313 classified islet cells, 328,225 were β- cells.

Using these cell annotations, we analyzed β-cells along T1D disease stages. As expected, there was a gradual loss of β-cells from pancreatic islets with advancing disease stage^1^ (Figure 1E). In non-diabetic donors, β-cells constituted on average 50% of islet cells, whereas donors with long-standing T1D had almost no β-cells (Figure 1E). We found considerable inter-donor variation within the mAAb+ and Onset T1D groups, indicating that the extent of β-cell loss was donor-specific. Younger age and longer time since disease onset were associated with lower β-cell fractions for Onset donors (not shown). Loss of β-cells was accompanied by an expected increase in the α-cell fractions (Figure S3B), and a potential loss in ι-cells in Onset donors. γ and δ-cells did not show major changes in abundance across disease stages (Figure S3B).

Analysis of key β-cell lineage, functional and stress markers across disease stages prior to LD T1D donors (due to their near-total loss of these cells) revealed a downregulation of several β-cell and islet-lineage markers as the disease progressed (Figure 1F, Figure S3C). This trend was most pronounced in β-cell hormones, with significant reductions (*P*<0.002) in IAPP, ProINS, and C- peptide observed in Onset T1D cases compared to controls (Figure S3C, Figure 1F). Notably, we detected significant loss of IAPP expression at the mAAb+ stage compared to controls (*P*=5*10^-3^). Slight non-significant losses of co-secreted INS and its precursor ProINS were observed at the mAAb+ stage compared to controls (both *P*≥0.8) implicating IAPP as a major marker of early disease progression. There was upregulation of HLA-ABC at the mAAb+ stage compared to the controls (*P*=0.08) and significant hyperexpression of HLA-ABC in Onset T1D donors compared to controls (*P*=9*10^-9^) (Figure 1F). Contrary to expectations^26,34,35^, we did not observe significant upregulation of ER-stress markers (ERN1, WFS-1, XBP1), inflammation markers (NFκB), or IFN-responsive markers (MX1, ADAR) across disease stages (Figure S3D). Only ADAR tended toward upregulation in Onset donors, indicating an IFN-response in clinical disease (Figure S3D). The TIGIT/CD226 receptor CD155 as well as the lowly-expressed redox signaling marker TXNIP were significantly downregulated in Onset donors^35,36^ (Figure S3D). In total, 24 out of 28 markers had lower protein levels in Onset than in control donors. These data suggest that the functions of remaining β-cells degrade as T1D progresses, including in clinically targeted pathways such as TXNIP^37^. Finally, we tested for age-associated differences in β-cell marker expression. We did not observe significant differences for any β-cell markers but did observe a trend toward higher levels of expression of IFN-responsive markers and lower levels of β-cell hormones in younger versus older subjects in Onset donors, suggesting more aggressive and IFN-driven disease in subjects younger than 13 years (Figure S3E).

Different islets from an individual may progress through T1D at different rates, potentially obscuring patterns of disease progression. Therefore, we inferred the temporal sequence of changes in the β-cells during disease progression using a pseudotime analysis^6^. We fit principal curves to diffusion map-embedded, single β-cell expression profiles using *slingshot*^38^. Even without specification of cluster labels or start and end nodes, *slingshot* detected a single lineage (Figure 1G), which indicates that T1D progression can be modeled along one main trajectory, despite donor heterogeneity. The predicted pseudotime started at control donors, progressed with sAAb+ and mAAb+ as mixed samples between the two endpoints and ended with Onset T1D donors (Figure S3F, S3G). More specifically, it ended in β-cells localized in insulitic islets, i.e. immune infiltrated islets with at least six T cells immediately adjacent to or within the islets^39^ (Figure S3H). The average β-cell pseudotime per donor thus increased significantly over disease progression (Figure 1H), and was well correlated to known features of T1D progression such as donor HbA1c status (*r*=0.64, *P*=10^-7^), β-cell fraction (*r*=-0.74, *P*=5*10^-14^), and disease stage (*r*=0.75, *P*=2*10^-14^), confirming the biological relevance of the inferred trajectory.

Analyses of β-cell marker correlations to pseudotime showed that IAPP downregulation and HLA-ABC upregulation were the β-cell marker trajectories most strongly associated with disease progression (Figure 1I; Figure S3I, both *P*<10^-16^). Downregulation of ProINS was on average observed only in Onset cases, whereas IAPP was significantly downregulated in some mAAb+ islets, which apparently were IAPP+INS- (Figure 1F, I, J, Figure S3I). As in our cross-sectional analysis, we observed gradual, but slighter, downregulations in other lineage β-cell markers such as PDX1, NKX6.1, and INS over pseudotime, but no significant changes in ER stress, inflammation, or IFN-responsive markers (Figure 1I). Our data therefore suggest that ProINS and INS loss is *preceded* by loss of IAPP, despite their known co-secretion^40^.

In summary, starting in the mAAb+ stage, we observed a gradual downregulation of β-cell lineage markers, with strong downregulation of IAPP, and progressive upregulation of HLA-ABC across pseudotime with loss of β-cell hormones prior to β-cell death. Again, we observed downregulation in 24 out of 28 measured β-cell markers, including mild and significant downregulation across ER-stress and other functional markers, suggesting degradation of β-cell function along disease progression.

### MHC-I, MHC-II and other IFN-response markers are upregulated in islet cells alongside immune cell infiltration

Islet immune infiltration as well as β-cell MHC-I hyperexpression are main hallmarks of T1D^3^. A controversial question has been whether MHC-II is hyper-expressed on β-cells during T1D development^41,23^, given the potential implications for direct antigen-presentation from β-cells to CD4^+^ T cells and that initial MHC-II observations on β-cells could not be reproduced. MHC-II hyperexpression was recently reported in pancreatic ductal cells^30^ as well as in β-cells^42,43^ of T1D patients. Highly multiplexed imaging techniques are needed to establish HLA-DR (MHC-II) expression not only in β-cells but also other islet cells, and further to exclude spillover or false annotation. Using our extensive immune panel, we therefore probed for MHC-II hyperexpression in our cohort and also tested for co-expression with MHC-I.

We observed increases in single-cell expression of both HLA-ABC (MHC-I) and HLA-DR (MHC-II) in multiple endocrine and exocrine cell types across T1D stages with significantly higher expression in Onset T1D relative to control donors and with stronger upregulation of HLA-ABC than HLA-DR (Figure 2A-C). Notably, we observed significant MHC-I upregulation in α-cells in mAAb+ donors, demonstrating that islet-cell perturbations extend to non β-cells prior to clinical onset (Figure 2A). In islets, antigen-presenting cells (APCs) and inflamed endothelium expressed HLA-DR at high levels, but we also clearly detected HLA-DR^+^ β-cells without potential spillover from major APC, phagocyte or endothelial markers in inflamed islets (e.g., VIM, CD204, CD20, CD11c, CAV1) (Figure 2C). Further, we also observed other HLA-DR+ endocrine and exocrine cells, even though at lower HLA-DR intensity than in β-cells (Figure 2B,2C). We observed significant co-expression of HLA-ABC in various islet cell types (for α- and β-cells, Spearman *r*=0.78-0.97, *P* <10^-16^; for β-cells and acinar cells, *r*=0.49-0.73, *P* <10^-16^), and co-expression increased with disease stage (Figure S4A). This suggests that a pro-inflammatory microenvironment upregulates islet and islet-proximal HLA-ABC protein levels. Levels of HLA-DR were not as strongly correlated (e.g., for α- and β-cells *r*=0.77-0.81, *P* <10^-16^) (Figure S4B). To determine the (pseudo)temporal sequence of HLA-DR and HLA-ABC upregulation, we aligned expression of these markers in β-cells along pseudotime. This revealed that HLA-ABC upregulation in these cells occurs prior to upregulation of HLA-DR (Figure 2D). HLA-DR upregulation was observed only in islets with strong HLA-ABC hyperexpression and only in late pseudotime (Figure S4C).

**Figure 2:**
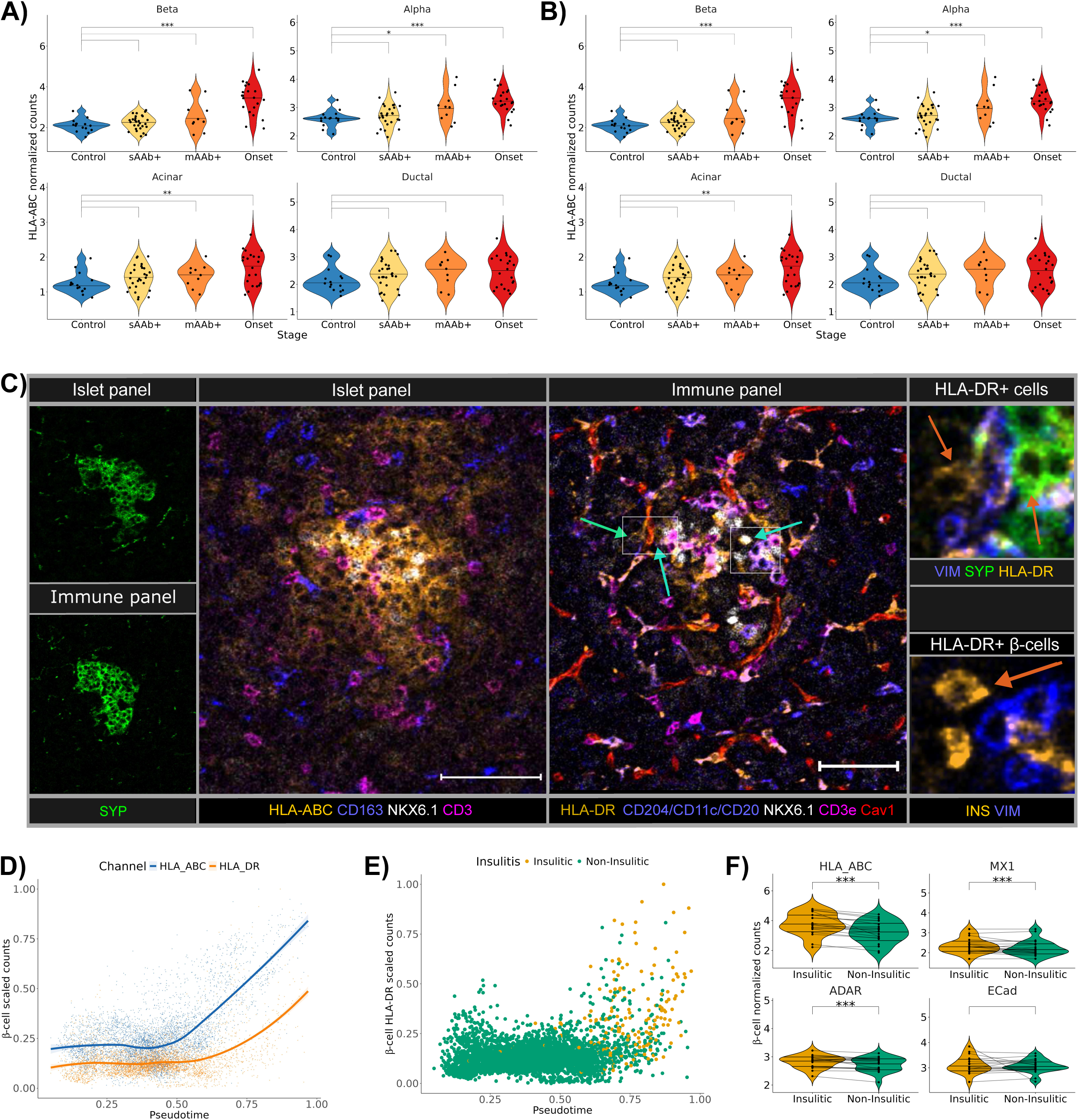
Interferon-dependent MHC-II upregulation in β-cells and further endocrine and exocrine cells. **A and B:** Violin plots of average (A) HLA-ABC and (B) HLA-DR single-cell expression in the indicated cell types by disease stage. Each dot indicates data from one donor of marker expression averaged across ROIs. HLA-ABC was analyzed in experiments with the islet antibody panel and HLA-DR with the immune antibody panel. Linear mixed effects models were fit to determine significance of differential expression for each stage in comparison to non-diabetic controls. **C:** Representative images of two consecutive sections stained with the islet and the immune antibody panels. Low magnification images (left) mark islets with SYP in both panels. Markers shown in the high-magnification images (right) are indicated. Arrows indicate HLA-DR^+^ β-cells (orange) that express NKX6.1 (white) but not APC markers (purple), or other HLA-DR^+^ endocrine and exocrine cells. Arrow in the magnified upper inset indicates HLA-DR+ endocrine (SYP+) cells, and HLA-DR+ exocrine cells (SYP-). Arrow in the magnified lower inset indicates INS^+^ β-cell (orange), negative for VIM (a mesenchymal cell marker; blue). Scale bar is 75 µm. **D:** HLA-ABC and HLA-DR expression in β-cells plotted over pseudotime. Each dot is the average expression per ROI; values are min-max scaled. Smoothing lines were fitted with locally weighted scatterplot smoothing (LOWESS) with the shaded areas indicating the confidence interval of 95%. **E:** HLA-DR expression in β-cells plotted along pseudotime. Each dot is the average expression per ROI with color indicating insulitic and non-insulitic ROIs. **F:** Violin plots of mean expression levels of the indicated markers between ROIs with or without insulitic islets. Only β- cells from Onset T1D donors were considered. Mean β-cell expression levels of 97 insulitic ROIs across 15 donors were compared against mean β-cell expression levels of 795 non-insulitic ROIs across 21 donors. Linear mixed effects models were fit to determine significance. Each dot indicates the average expression per sample across insulitic or non-insulitic ROIs. Lines connect insulitic and non-insulitic ROIs from the same donor. Tests: ^∗^ *P* <0.05; ^∗∗^ *P* < 0.01; *^∗∗∗^ P* <0.001 for all comparisons. Significance was determined for each stage by testing for differential abundance against controls using *edgeR* and by testing for differential expression against controls by fitting a linear mixed-effects model.

More specifically, upregulated HLA-DR expression in β-cells was correlated with abundance of myeloid and especially T cells in mAAb+ and Onset T1D donors (Figure S4D) and was mostly confined to insulitic islets (Figure 2E). In contrast, HLA-ABC upregulation was not confined to insulitic islets (Figure S4E). β-cells in insulitic islets had higher levels not only of HLA-DR and HLA-ABC relative to non-insulitic islets, but also of IFN-responsive markers MX1 and ADAR (Figure 2F). While MX1 was upregulated in insulitic islets, MX1 was *not* upregulated in islets of Onset donors in comparison to control donors (Figure S3D), suggesting an IFN-driven β-cell state specifically associated with T cell infiltration. Importantly, as for HLA-DR, MX1 and ADAR were also upregulated in other islet cells, albeit at lower effect size than β-cells (Figure S4F). Finally, we also observed upregulation of adhesion marker CD54 (also known as ICAM-1) in β-cells in insulitic islets (Figure S4G), and in endothelial cells as expected^44^ (not shown), but no significant differences were detected in ER-stress markers or other functional markers of β-cells between insulitic and non-insulitic islets (Figure S4H). In sum, these data suggest that in islets, MHC-I is upregulated first, coinciding with protein downregulation in most lineage and functional β-cell markers, followed by upregulation in MHC-II and other IFN-responsive markers during T cell infiltration. Importantly, these changes are observed in β-cells, but also in other islet and islet-proximal endocrine and exocrine cells, suggesting that they are not β-cell specific processes, *per se*.

### Immune cell types change in density during disease progression

To study the evolution of the islet and islet-proximal exocrine immune compartment during T1D progression, we used the immune panel data for immune phenotyping, differential abundance analysis, immune infiltration analysis, and alignment of these data with β-cell pseudotime. Identified immune cell types of interest were then sub-clustered and analyzed in detail (Figure 3A). As with the islet-focused antibody panel, we implemented a multi-step annotation approach using clustering of cellular lineage marker expression profiles, manual annotation of clusters, and automated assignment based on the consensus of four annotation approaches. We first separated cell types into immune and non-immune cell types and classified non-immune cell types as acinar, ductal, mesenchymal, nerves, endothelial, smooth-muscle, or ‘other’. Next, immune cells were annotated as myeloid cells, neutrophils, natural killer (NK) cells, B cells, CD4^+^ helper T (T-CD4) cells, CD8^+^ cytotoxic T (T-CD8) cells, double-negative T (T-DN) cells (CD3e^+^CD4^-^ CD8^-^, which likely include natural killer T cells and γδ T cells), and CD303^+^/VIM^+^ cells (Figure 3B; Figure S5A). CD303^+^/VIM^+^ cells are likely a fibroblast subset involved in the fibrotic process after β-cell destruction or potentially an atypical plasmacytoid DC (pDC; CD303^+^HLA-DR^-^) (Figure S5A). In total, about 10.5 million cells were annotated, and approximately 1.2 million were immune cells (Figure 3B).

**Figure 3:**
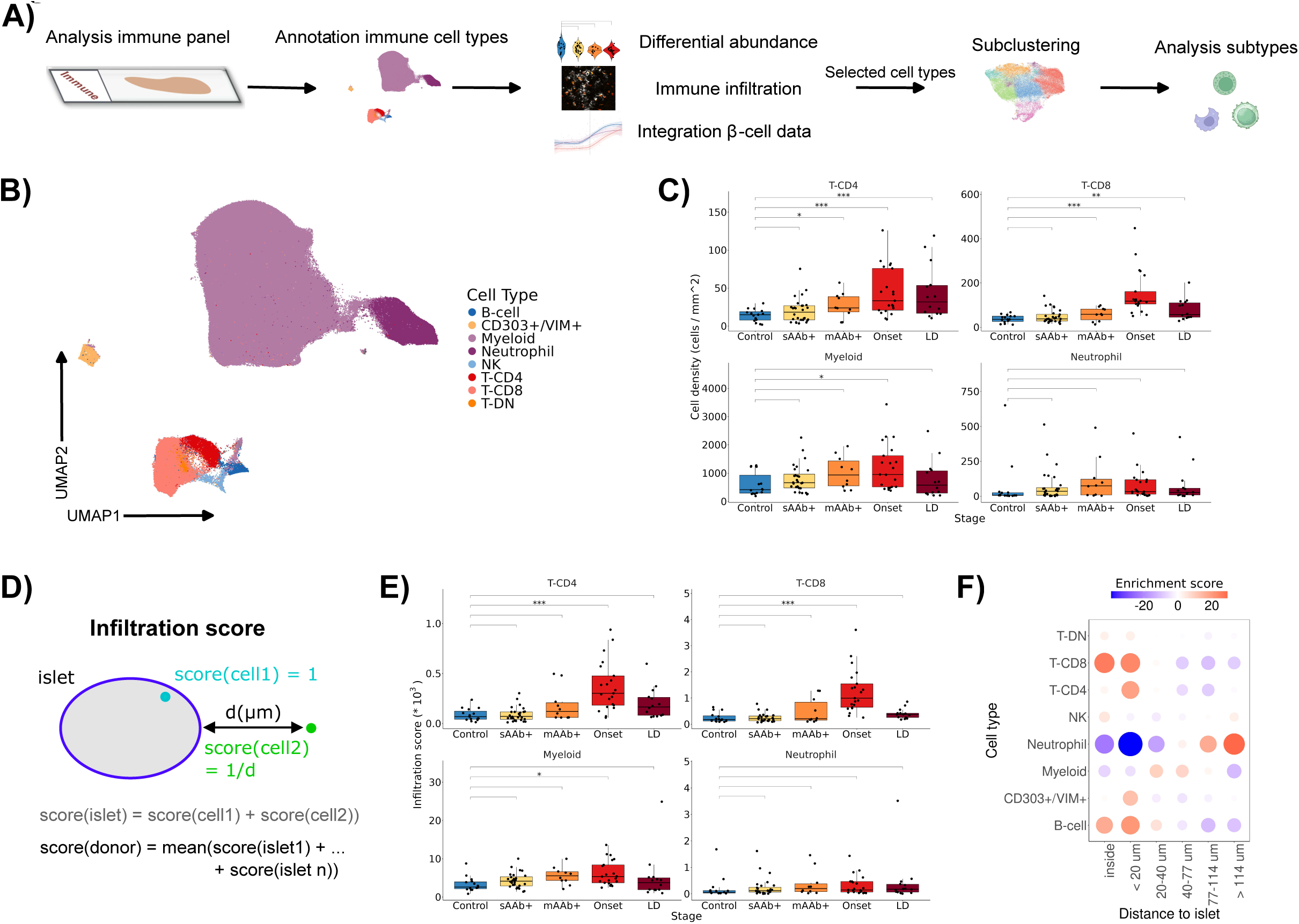
Immune cell type dynamics across T1D disease stages. **A:** Data analysis with the immune panel. **B:** UMAP of immune cell expression profiles, colored by cell type. **C:** Mean densities of the indicated cell types in control and diseased samples. Each dot indicates data from one donor with mean cell density averaged across ROIs. Significance was determined for each stage by testing for differential abundance against controls using *edgeR*. **D:** Sketch of the developed infiltration score. The infiltration score is additionally normalized by the total number of cells in the acquired ROI. **E:** Mean infiltration scores of the indicated cell types in control and diseased samples. Each dot indicates data from one donor averaged across ROIs. Significance was determined for each stage by a linear mixed-effects model and comparison against non-diabetic controls. **F:** Enrichment score of immune cell type enrichments in distance bins relative to the islet edge. Red indicates enrichment, blue indicates absence. Enrichment score values are χ^2^-residuals. Tests: ^∗^ *P* <0.05; ^∗∗^ *P* <0.01; ^∗∗∗^ *P* <0.001 for all comparisons. Significance was determined for each stage by testing for differential abundance against controls using *edgeR* and by testing for differential infiltration score against controls by fitting a linear mixed-effects model.

We quantified immune cell density along T1D progression to identify disease-relevant immune cell types. Myeloid, B, and T cells displayed gradual pancreas infiltration beginning in early disease stages, with significant increases over control samples in Onset donors for all immune cell types (Figure 3C, Figure S5B). T-CD4 density was increased significantly at the mAAb+ stage relative to the controls (*P*=0.04). B cell density was variable between donors with significantly higher fractions in younger T1D donors than in those ≥13 years of age (Figure S5C), as expected based on previous reports^45^. Neutrophil densities were variable, with outlier cases within each donor group showing considerable neutrophil presence (Figure 3C). CD303^+^/VIM^+^ cells were present at significantly higher density in LD T1D samples than in other groups (Figure S5B). There were no significant differences in densities for NK or T-DN cells along T1D progression (Figure S5B).

Both increase in abundance of immune cells within the pancreas and attraction to islets are likely relevant for T1D progression. Therefore, we developed an “infiltration score” that combines distance to islet edge and cell abundance in the measured ROIs (Figure 3D). We observed a highly significant increase in infiltration score for T cells, and to some extent for B and myeloid cells, in Onset T1D donors compared to controls (Figure 3E, Figure S5D). Along pseudotime, we observed co-infiltration of T-CD4 and T-CD8 cells along pseudotime, including into islets of mAAb+ donors (R=0.68, *P*<10^-16^), with myeloid cells potentially preceding this co-infiltration (*P*=0.2) (Figure S5E). Other cell types such as neutrophils showed no changes in infiltration score along T1D disease stages, with neutrophils displaying specific islet avoidance (Figure 3E,F, Figure S5D). Based on these results, we focused our subsequent analyses on the major islet-infiltrating immune cells, i.e. T-CD4, T-CD8, and myeloid cells.

### PD1^+^ CD4^+^ T cells invade the pancreas early in T1D progression and exhausted-like eFector CD8^+^ T cells dominate islet infiltration in mAAb+ and T1D donors

T cell targeting therapies have shown partial therapeutic success prior to and after clinical onset of T1D^7^. However, while T cell states have been comprehensively evaluated in peripheral blood, they remain incompletely characterized in human pancreata, especially from AAb+ patients. We thus analyzed potential T-CD4 and T-CD8 states in our cohort.

We first clustered around 27,000 T-CD4 cells using Leiden clustering and assigned them to eleven T-CD4 subtypes (Figure 4A). Clusters included activated and minimally activated CD45RO^+^CD45RA^-^CD27^-^ effector memory-like cells (T-EM-like act., T-EM-like low act.), CD45RO^+^CD45RA^-^CD27^+^ central memory-like cells (T-CM-like act., T-CM-like low act.), PD1^+^ T- CD4 memory cells (PD1+ act., PD1+ low act.), PD1^low^ cells (PD1^low^ act., PD1^low^ low act.), as well as two regulatory phenotypes, CD73^+^ T-CD4 cells and regulatory T cells (T-regs). We also labeled T-CD4 cells with ambiguous expression profiles and an undefined T-CD4 cluster (T-CD4 Other). Next, we clustered around 72,000 T-CD8 cells and assigned them to eight T-CD8 cell subtypes (Figure 4B): CD45RA^+^CD27^+^CD57^-^ naive T-CD8 cells, CD45RA^+^CD27^low^CD57^+^ effector memory cells re-expressing CD45RA T cells (T-EMRA), activated and marginally activated CD45RO^+^CD103^+^ tissue-resident memory cells (T-RM act., T-RM low act.), CD45RO^+^CD27^mid^ effector memory/central memory-like cells (T-EM/CM-like), GranB^+^ cytotoxic T-CD8 cells, an undefined set of T-CD8 cells (T-CD8 Other), and a T-CD8 memory cell subtype that expressed high-levels of markers of exhaustion (PD1, TIM-3), cytotoxicity (GranB), survival (CD27), and metabolism (e.g., citrate synthase (CS)), suggesting an exhausted-like phenotype with some cytotoxic effector functions (T-ex^e^^7^).

**Figure 4:**
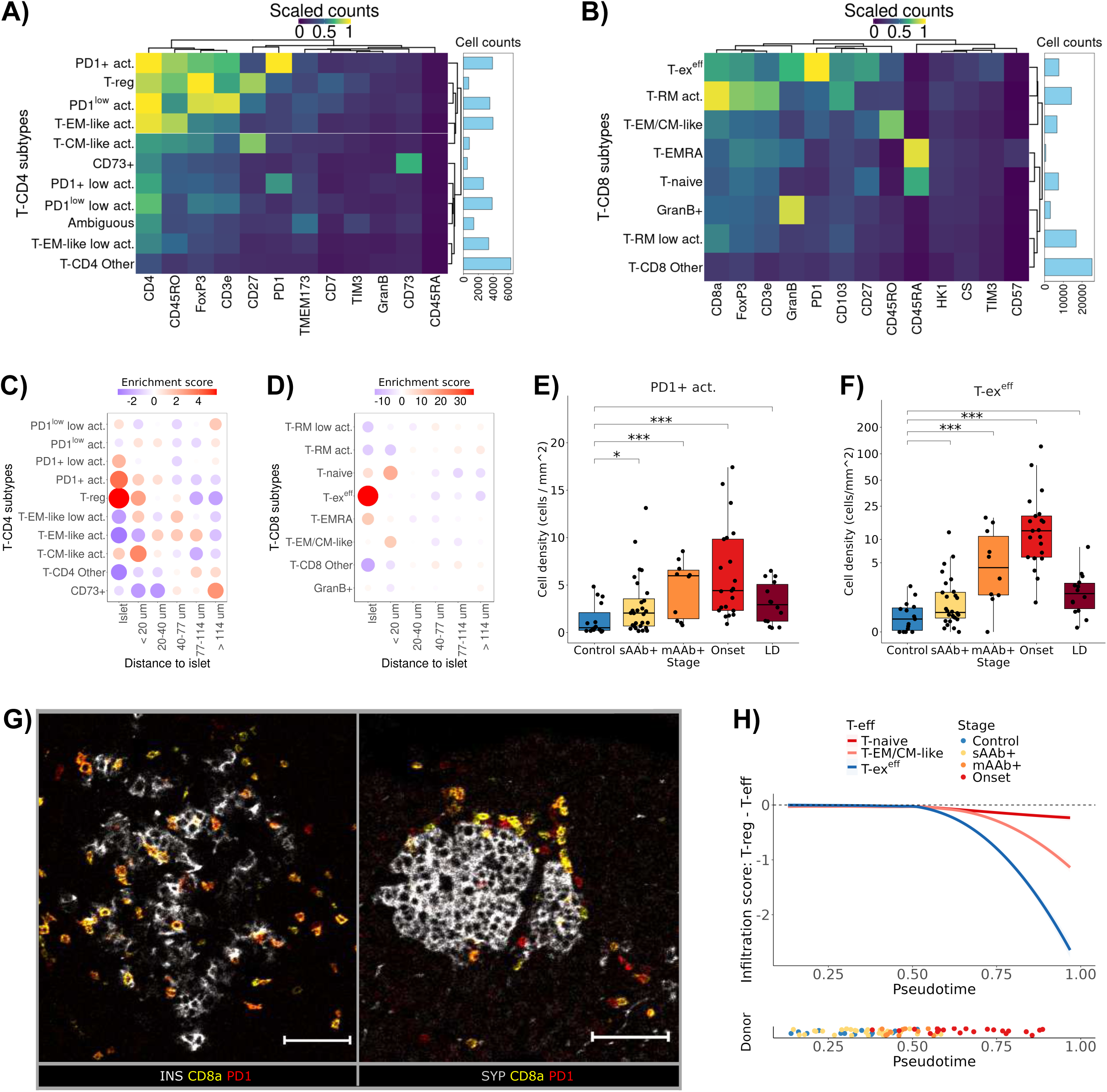
Exhausted-like T cells dominate islet infiltration in mAAb+ and T1D donors. **A:** Lineage marker expression levels in the 11 annotated T-CD4 cell subtypes. The barplot to the right indicates their total cell counts. **B:** Lineage marker expression in the eight annotated T-CD8 cell subtypes. The barplots to the right indicate total cell counts. **C and D:** Enrichment scores of T- CD4 **(C)** and T-CD8 cell subtypes **(D)** in distance bins relative to the islet edge. Red indicates enrichment, blue indicates absence. Matrix elements are the χ^2^ residuals. **E and F:** Boxplots of mean densities of PD1+ act. cells **(E)** and T-ex^e^^7^ cells **(F)** in control and diseased tissues. Each dot indicates the mean cell density per sample averaged across ROIs. **G:** Representative images of islet (left) and peri-islet enrichment (right) of exhausted-like T cells from an Onset donor and mAAb+ donor. Left: Staining highlighting β-cell (INS; gray) interaction with T-ex^e^^7^ cells (orange), i.e. cells costained for CD8a (yellow) and PD1 (red). PD1^+^CD8a^-^ cells are PD1+ act. cells (red). Right: Staining of a mAAb+ donor sample stained for islet cell marker (SYP; gray), CD8a (yellow), and PD1 (red). PD1^+^CD8a^+^ (orange) cells are T-ex^e7^ cells. PD1^+^CD8a^-^ cells (red) are PD1+ act. cells. The scale bar is 75 µm. **H:** Infiltration scores of distinct T-CD8 effector cells (T-eb) along pseudotime per ROI. Infiltration scores were subtracted against infiltration scores of T-regs. Negative values thus indicate higher presence of T-eb than T-reg cells in islets. Regression lines were fitted with LOWESS, with the shaded areas indicating the confidence interval of 95%. Average pseudotime per donor is indicated by the x-axis position of the labeled dots in the lower panel. Tests: ^∗^*P* <0.05; ^∗∗^*P* <0.01; ^∗∗∗^*P* <0.001 for all comparisons. Significance was determined for each stage by testing for differential abundance against controls using *edgeR* and by testing for differential expression against controls by fitting a linear mixed-effects model.

Next, we analyzed islet-enrichment and abundance of T cell subtypes across progression. For T- CD4 cells, we observed islet-enrichment of T-regs (*P*=5*10^-7^) and PD1+ act. cells (*P*=4*10^-4^) (Figure 4C), and for T-CD8 cells, we observed strong islet-enrichment of T-ex^e^^7^ cells (*P*<10^-16^) (Figure 4D). In an analysis across disease stages, the islet-enriched PD1+ subtypes (i.e., T-ex^e^^7^ and PD1+ act. cells), demonstrated significantly greater abundance in mAAb+ and Onset donors compared to controls (all *P*<10^-5^) (Figure 4E, 4F), with PD1+ act. cell abundance also higher already in sAAb+ donors (*P*=0.047, Figure 4E). Also enriched in mAAb+ samples were T-regs, GranB+ cytotoxic T-CD8s and T-RM act. cells (Figure S6A,B). However, GranB+ cytotoxic T-CD8s and T-RM act. cells were not enriched in islets (Figure S4D), suggesting limited cytotoxic effects at this stage of disease.

PD1+ act. and T-ex^e^^7^ cells significantly infiltrated islets in Onset donors and a subset of mAAb+ donors, but not in sAAb+ donors, compared to controls (Figure S6C). In sAAb+ donors, PD1+ act. cells were mostly confined to the islet-proximal pancreas (Figure S6D). Notably, infiltration scores of PD1+ act. and T-ex^e^^7^ cells subtypes displayed the highest association to β-cell MHC-I and MHC-II expression of all T cell subtypes along pseudotime (all R>0.85, *P*<10^-14^), with infiltration following MHC-I and coinciding with MHC-II expression on β-cells (Figure S6E). With disease progression, we observed upregulation of exhaustion-markers PD1 and TIM-3 for both subtypes, as well as tissue-residency marker CD103 for T-ex^e^^7^ cells, which might also signal exhaustion^46^, but not of the cytotoxicity marker GranB (Figure S6F, G), together suggesting exhaustion of these subtypes within the islet microenvironment.

Next, we analyzed the spatial niches of T cell subtypes to determine if these were reflective of diabetogenic processes. We identified the 10 nearest neighbors of all cells, clustered the resulting cell populations into 60 cellular neighborhoods (CNs)^47^, and then manually annotated them by their enriched cell types, localization, and disease stage association (Figure S7A). Similar CNs were aggregated into 22 CNs. This analysis showed stronger enrichment of PD1+ act. cells and T-ex^e^^7^ cells in β-cell enriched CNs (‘Beta’) than in other islet-cell CNs (Figure S7B, C), with especially T-ex^e^^7^ cells often in direct contact with β-cells (Figure 4G). We also observed enrichment of both subtypes at the islet-edge in mAAb+ donors (‘Islet-edge mAAb+’), which likely captures peri-islet accumulation in mAAb+ donors prior to overt islet infiltration (Figure S7B, C; Figure 4G).

In Onset donors, we observed enrichment for most non-exhausted T cell subtypes (Figure S6A, B). Both infiltration scores and enrichment in T cell rich CNs (‘T-CD4 > T-CD8’, ‘T-CD8 > T-CD4’) suggested key roles for T-CD4 PD1^low^ act. cells, which are likely early effector cells (Figure S7B), as well as T-naïve and T-CM/EM-like cells (Figure S7C). These non-exhausted T cell subtypes appeared to follow accumulation of exhausted-like T cells into islets, apparently outcompeting T-regs in islets of Onset donors (Figure 4H; Figure S7D).

In sum, this indicates the PD1+ T cell subtypes as critical indicators of early disease, with progressive shifts from islet-proximal tissue sites in AAb+ donors, to overt islet infiltration in Onset donors. Their specific β-cell directed infiltration is strongly linked to islet inflammation, especially MHC-II. While the islet micro-environment appears to limit cytotoxicity of these cells by further exhaustion upon infiltration, with potentially also T-regs playing a role, expression profiles of T-ex^e^^7^ cells suggest remaining cytotoxicity and thus contribution to β-cell demise. Afterwards, non-exhausted T cells infiltrate islets, this could suggest the loss of a T cell exhaustion checkpoint after MHC-II linked IFN footprints in islets.

### Peri-islet macrophages are M1 polarized

In addition to T cells, myeloid cells are the main immune cells that infiltrate islets during T1D progression (Figure 3). Pancreatic myeloid cells were shown to be important for T1D pathogenesis in mouse models^16^, but the role of myeloid cells has been scarcely studied in humans^22^. We therefore analyzed contributions of myeloid cells to disease progression.

We subclustered the approximately 800,000 myeloid cells in our dataset and then used a combination of lineage marker expression, activation states, and spatial location within the pancreas to annotate nine distinct myeloid subtypes (Figure 5A). Clusters included activated and minimally activated exocrine macrophages (Exocrine act.; Exocrine low act.), and (peri)-islet enriched M1/M2-like macrophages (M1/M2-like act., M1/M2-like low act.). These islet-resident macrophages were distinguished by their expression levels of both classical M1 and M2 markers (HLA-DR, CD163, CD206) (Figure 5A, 5B). Further, we annotated conventional DCs (cDCs), characterized by high levels of TIM-3, HLA-DR, and CD11c and low levels of macrophage markers CD163 and CD206. These cells were rare (∼1100 cells) and have, to our knowledge, not been defined in human multiplexed T1D imaging studies so far. We annotated further clusters as CD54^+^ macrophages, Exocrine MPO^+^ macrophages, CD11c^+^ macrophages, and cells with non-myeloid-specific expression profiles as ‘Ambiguous’.

**Figure 5:**
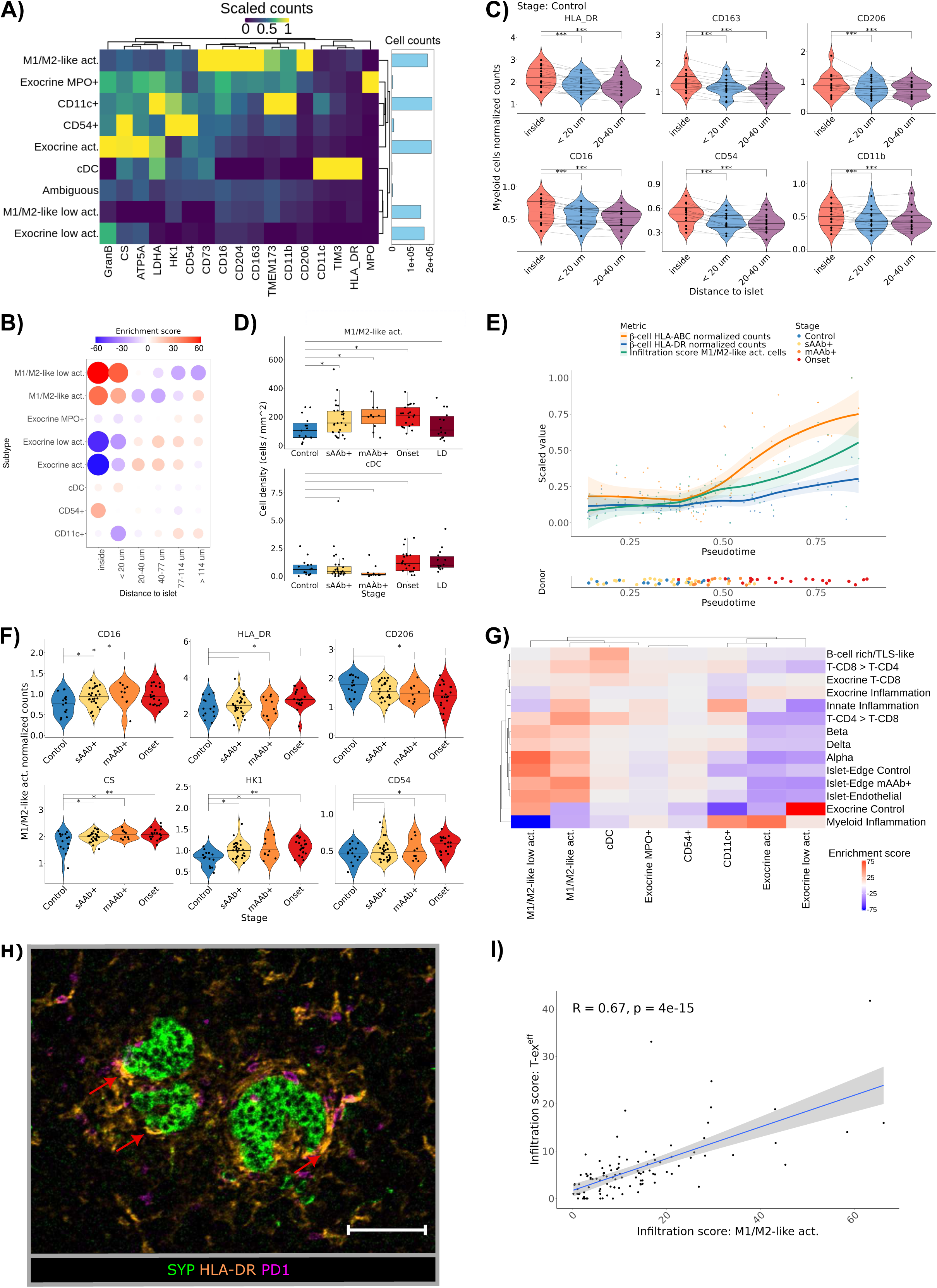
Activated M1/M-like macrophages and cDCs interact with lymphocytes at the islet periphery and within islets. **A:** Lineage marker expression in the nine annotated myeloid cell subtypes. The barplot to the right indicates total cell counts. **B:** Enrichment scores of myeloid cell subtypes in distance bins relative to the islet edge. Red indicates enrichment, blue indicates absence. Matrix elements are χ^2^ residuals. **C:** Expression of key functional and lineage markers of islet and peri-islet myeloid cells from 15 non-diabetic donors. Each dot indicates data from one donor with mean expression levels averaged across ROIs. **D:** Boxplots of mean densities of the indicated myeloid cell subtypes in control and diseased tissue. Each dot indicates data from one donor with cell density per sample averaged across ROIs. **E:** Scaled expression of HLA-ABC and HLA-DR in β-cells and infiltration scores of M1/M2-like act. macrophages visualized along the inferred β-cell pseudotime trajectory. Each dot is the mean value per donor. **F:** Levels of key myeloid and metabolic markers in activated M1/M2 macrophages, compared across disease stages. Each dot indicates the average marker expression across ROIs for each sample. **G:** χ^2^-test residuals for enrichment of the indicated myeloid cell subtypes in informative CNs. Red indicates enrichment, blue indicates absence. **H:** Image of ROI from mAAb+ donor with significant peri-islet immune infiltration. Red arrows indicate HLA-DR^high^ cDCs (orange) at the SYP^+^ islet edge (green) in close contact with PD1+ T cells (magenta). HLA-DR^+^ M1/M2-like act. macrophages (orange) are most peri-islet macrophages (without arrows). Scale bar is 75 µm. **I:** Infiltration scores of T-ex^e^^7^ cells and M1/M2-like act. cells in insulitic islets. Infiltration scores are unnormalized. Tests: ^∗^ *P* <0.05; ^∗∗^ *P* <0.01; ^∗∗∗^ *P* <0.001 for all comparisons. Significance was determined for each stage by testing for differential abundance against controls using *edgeR* and by testing for differential expression against controls by fitting a linear mixed-effects model.

Almost all subtypes showed a gradual increase in abundance during disease progression (Figure S8A). For example, Exocrine MPO^+^, Exocrine act., and CD54^+^ macrophages demonstrated significantly higher abundances in Onset donors than in donors at earlier stages of disease (Figure S8A). This highlights the importance of immune cell composition within the peri-islet exocrine niche for T1D progression. Several key myeloid markers (CD163, CD206, CD16, HLA-DR, CD54, CD11b) were more strongly expressed in islet-resident myeloid cells than in myeloid cells outside islets in all disease stages (Figure S8B), including in non-diabetic controls (Figure 5C), suggesting also that islet-resident myeloid cells are activated and are relevant in both homeostasis and disease.

Given their roles in peripheral immunity/tolerance, M1/M2-like act. macrophages and cDCs are likely critical in T1D^48^. We detected increased frequencies of M1/M2-like act. macrophages in the sAAb+ stage compared to controls, with some inter-patient variability (Figure 5D). In contrast, cDCs did not show an obvious increase in frequency over disease progression (Figure 5D). We did observe a shift of both cDCs and M1/M2-like act. macrophages to the islet edge in mAAb+ and Onset donors relative to controls, evident in their distances to islets and their infiltration scores (Figure S8C, D). The M1/M2-like act. macrophage infiltration score followed the same trajectory over pseudotime as HLA-ABC expression in β-cells, especially during initial HLA-ABC upregulation, and showed cross-correlation to HLA-DR in later pseudotime (Figure 5E). This suggests that infiltration of these macrophages is associated with both early and late islet inflammation. Marker expression in M1/M2-like act. macrophages differed during disease progression, with gradual upregulation of metabolism markers (e.g., CS, HK1), significant upregulation of IFN-dependent pro-inflammatory markers (CD54, HLA-DR) and higher expression of monocyte markers (CD16) over the disease course (Figure 5F). Interestingly, some of these expression changes were observed early in the disease (i.e., CS, CD16, and HK1 differed from control levels in the sAAb+ stage). We also observed downregulation of classical M2- markers, such as CD206, over the disease course (Figure 5F, Figure S8E). Differences in levels of other major myeloid markers were not significant (Figure S8F). We then investigated whether the phenotypes of these M1/M2-like act. macrophages change after the loss of β-cells by comparing islets with (ICI) and without (IDI) β-cells from the same Onset donors. We observed upregulation of classical M2-markers (i.e., CD163, CD206) and downregulation of activation and pro-inflammatory markers (e.g., CD54, HLA-DR) in M1/M2-like act. macrophages from IDIs compared to ICIs (Figure S9A, B). Similar results were observed when markers from the islet panel were analyzed (e.g., NFκB, CD155, MHC-I; Figure S9C). Overall, these data suggest that islet inflammation that occurs early in the disease course results in attraction, activation, and M1- polarization of islet-resident M1/M2-like act. macrophages. This likely also includes attraction of monocytes. After loss of β-cells, macrophages return to an M2-like phenotype.

Next, we analyzed enrichment of myeloid subtypes in CNs. Importantly, both M1/M2-like act. macrophages and cDCs were significantly enriched in T cell enriched CNs (‘T-CD8 > T-CD4’ or ‘T- CD4 > T-CD8’), suggesting specific myeloid-lymphocyte interactions with both T-CD4 and T-CD8 cells (Figure 5G). M1/M2-like act. macrophages were enriched around islet-endothelial cells and the islet-edge, spatially positioning them as key regulators of invading lymphocytes. Interestingly, they were especially enriched at the islet-edge in mAAb+ donors, just as exhausted-like T cells (Figure S7B,C), suggesting key interactions between myeloid cells and exhausted-like T cells in peri-islet staging areas prior to overt islet infiltration (Figure 5G). Indeed, we observed cDCs and M1/M2-like act. macrophages in direct contact with T cells, including exhausted-like T cells, at the islet-periphery (Figure 5H). Further, M1/M2-like act. macrophages and cDCs were enriched in insulitic islets (Figure S9D), where both cell types displayed significant upregulation of CD54 and HLA-DR (Figure S9E), and M1/M2-like act. macrophages displayed strong co-infiltration of insulitic islets with T-ex^e^^7^ cells (Figure 5I). This indicates IFN-dependent footprints in myeloid cells, linked to T-ex^e^^7^ cells. Further, this suggests cDC maturation and enhanced capacity for antigen-presentation and co-stimulation to infiltrating T-CD4 cells. Due to MHC-I hyperexpression on myeloid cells, as measured by the Islet panel (Figure S9C), cross-presentation capacity to T-CD8s might also be increased. In contrast, the high levels of TIM-3 that we detect on cDCs (Figure 5B) suggest a regulatory program that limits autoimmunity in the pancreas^49^. TIM-3 is slightly downregulated in cDCs in insulitic islets (Figure S9E). In sum, these data suggest that specific interactions between M1-polarized macrophages and cDCs with T cells, especially exhausted-like T cells, at the islet periphery and within islets can modulate autoimmunity.

### Identification of spatial, progression, and age-associated immune cell motifs

T1D severity varies across age, and the disease is known to be more precipitous in young patients^50^. To understand the potential basis of this effect, we sought to define potential age-associated immune cell motifs. First, to capture age-associated immune motifs in insulitis, we correlated infiltration scores of T-CD4, T-CD8 and myeloid cell subtypes in insulitic islets and compared abundance of the corresponding immune subtypes within these islets between younger (<13 years) and older (>13 years) donors. Second, to capture age-associated motifs affecting the homeostatic pancreatic immune niche, and potentially disease progression, we also compared abundance of immune cell subtypes in the entire sample across disease stages between younger and older donors. For both analyses, we excluded long-standing T1D donors due to the considerable time-lag between age of onset and age of donation.

Correlation of infiltration scores in insulitic islets, showed that cDCs and T-CD4 PD1+ act. cells formed a cluster with T-CD8 naïve and B-cells (Figure 6A; cluster 1); all these cells were more abundant in pancreata of younger donors across analyzed disease stages (Figure S10A). T-CD8 naïve and B-cells displayed especially strong co-infiltration (r=0.79, *P*=10^-14^), formed B-cell rich infiltrates or tertiary lymphoid structure (TLS)-like structures (Figure 6B), and were at higher abundance in insulitic infiltrates of younger donors in our cohort (Figure 6C), consistent with previous results^51,45^. This segregation across age was not complete, however, as examples of infiltrates with high B-cell content in older donors could also be found (Figure S10B). PD1+ act. cells and cDCs were not enriched in younger donor infiltrates (Figure 6C), and analysis revealed that co-infiltration, and thus likely interaction, was limited to few islets (Figure S10C).

**Figure 6:**
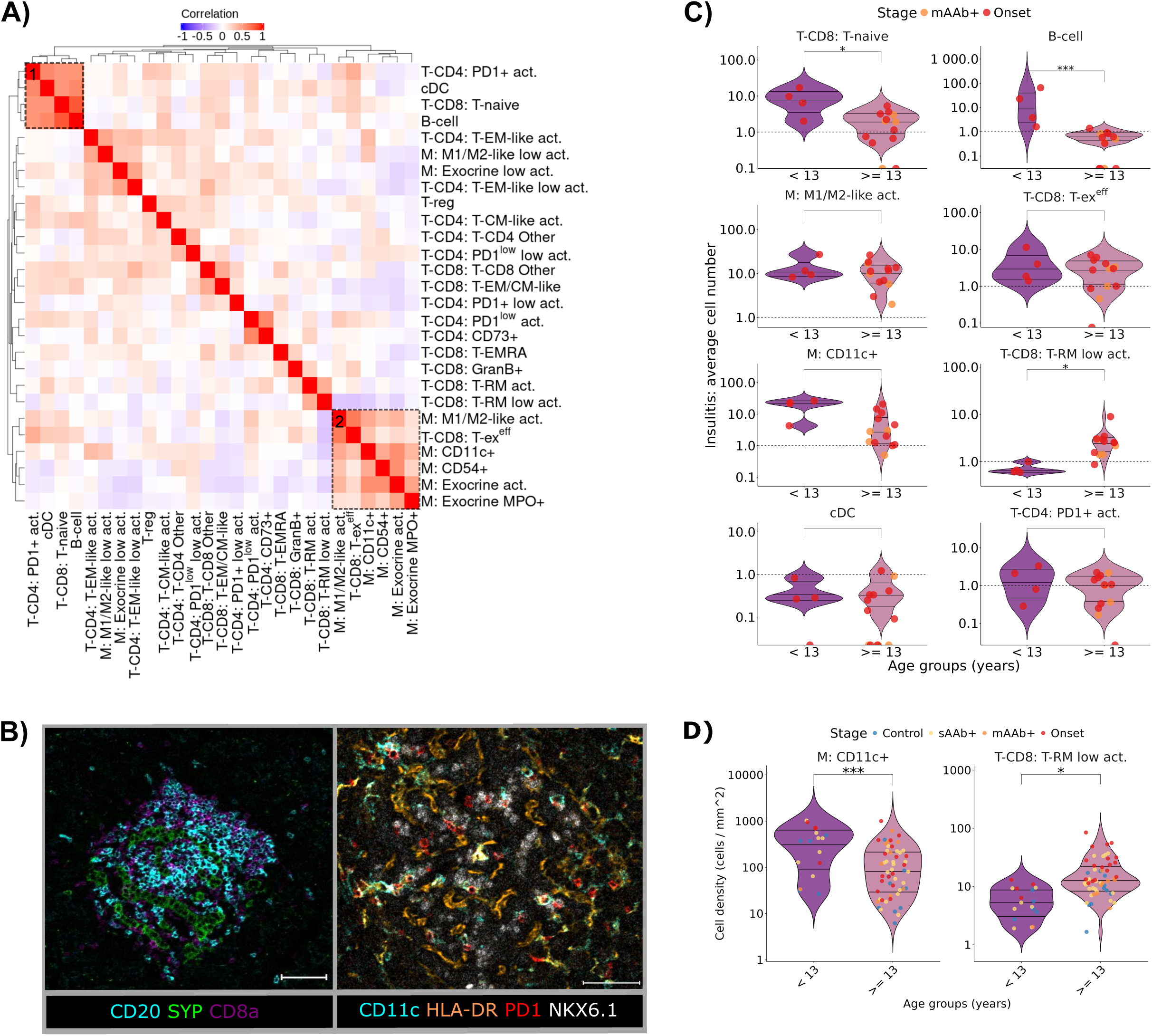
Age-associated immune cell phenotypes form insulitic clusters. **A:** Pearson correlation of infiltration scores of immune cell types (i.e., B cells and the indicated T-CD4, T-CD8 and myeloid (M) cell subtypes) from insulitic ROIs. Only islet and peri-islet cells (≤20 µm to the islet- edge) were considered. Numbers and brackets highlight different immune cell clusters. **B:** Micrographs of tissue from two recent-onset donors of <13 years of age. Left: Region of strong islet infiltration (SYP^+^) of both CD8a^+^ T cells and CD20^+^ B cells (cluster 1 in panel A). Right: Region of inflamed islet tissue area illustrating CD11c^+^HLA-DR^+^ macrophages in contact with PD1^+^ T cells (T-ex^e^^7^, PD1+ act.) (cluster 2 in panel A). NKX6.1 illustrates β-cells. **C:** Violin plots of mean immune cell numbers between insulitic infiltrates of younger (< 13 years) and older (≥13 years) donors. Each dot indicates the average immune cell number across insulitic islets per donor. Differential abundance was calculated using *edgeR*. **D:** Violin plots of mean density per sample between younger (< 13 years) and older (≥13 years) donors. Each dot is colored by disease stage and indicates the mean cell density per sample. Significance was calculated by a linear mixed- effects model, while adjusting for disease stage. Results for additional immune cell subtypes are shown in Figure S10A. Tests: ^∗^ *P* <0.05; ^∗∗^ *P* <0.01; ^∗∗∗^ *P* <0.001 for all comparisons.

A second motif in insulitic islets comprised multiple myeloid subtypes that clustered with exhausted-like effector T-CD8 (T-ex^e^^7^) cells, probably indicative of myeloid and T cell co-infiltration (Figure 6A; cluster 2; Figure 6B). This included M1/M2-like act. macrophages, as described above (Figure 5I), but also CD11c^+^ macrophages. Comparison of insulitic infiltrates between younger and older donors showed no strong differences for T-ex^e7^ cells and M1/M2-like act. macrophages (*P*>0.7), suggesting that this immune motif is present across age (Figure 6C). CD11c^+^ macrophages displayed only slight enrichment in insulitic islets of younger donors (*P*=0.07).

Interestingly, when we looked at abundance in the entire sample across disease stages, CD11c^+^ macrophages were strongly enriched in younger donors (Figure 6D), with a similar tendency visible also in controls (*P*=0.08). CD11c^+^ macrophages expressed the highest level of TMEM173 (STING) of all myeloid cells, suggesting that they are type 1 IFN producing cells^52^ (Figure 5B), and often expressed HLA-DR (Figure 5B, Figure 6B), suggesting increased capacity for antigen-presentation. These cells were enriched in spatial neighborhoods with high innate immune cell content (‘Innate inflammation’) (Figure 5G), which were more abundant in younger donors, as expected (Figure S10E), as well as in T cell enriched CNs (‘T-CD4>T-CD8’, ‘T-CD4<T-CD8’) (Figure 5G). In combination with their potential APC capacity and enrichment in insulitic infiltrates (Figure 6A), this also suggests interactions with T cells. Finally, older donors had higher abundances of marginally activated tissue-resident memory T-CD8 cells (T-RM low. act.) across disease stages than did younger donors (Figure 6D), again with similar trends in controls (*P*=0.08), as well as in insulitic infiltrates (Figure 6C). In comparison to other T-CD8 subtypes, T-RM low act. cells were enriched in exocrine tissue CNs found mainly in control donors (‘Exocrine Control’) (Figure S10D, Figure S7C). As expected, this CN was also enriched in older donors (Figure S10E). This suggests limited specificity of T-RM low act. cells for T1D.

In sum, when analyzing immune cell type abundance across disease stages and age, CD11c+ macrophages and marginally activated T-RMs were the main stratifying immune cells, suggesting that these differences might reflect evolution of the human pancreatic immune niche across age, with potential implications for T1D disease. When comparing insulitic infiltrates across age, M1- polarized macrophages and T-ex^e7^ cell motifs were observed across age, and B-cell rich / TLS-like structures enriched in younger donors, which might be associated with differential disease severity between younger and older donors.

## Discussion

T1D progression involves a complex interplay among endocrine, adaptive immune, and innate immune cells^3^. We report here a first of-its-kind cellular-resolution multiplexed imaging study of pancreatic samples from 88 donors across all stages of T1D, including sAAb+ and mAAb+, matched by age, gender, and BMI. Our analysis thus enabled interrogation of the human β-cell state and the pancreatic islet-immune interface during T1D progression at unprecedented scale.

Our single-cell IMC analysis tailored to investigate the β-cell state showed that human pancreatic β-cells are dysregulated prior to clinical disease onset. Most strikingly, we observed loss of the IAPP hormone in mAAb+ donors, a stage that precedes loss of co-secreted insulin^40^, leading to the observation of apparently IAPP- but INS+ islets. Non-detectable IAPP might be explained by loss of IAPP protein, but also by structural changes, especially since IAPP is known to be aggregation-prone^53^. Furthermore, it might be indicative of a more general stress response in β- cells^54^. In support of this notion, β-cells in diabetic Onset donors showed upregulation of MHC-I, and downregulation of 24 out of 28 β-cell lineage and functional markers in comparison to control donors. Our data offer several hypotheses of how these β-cell phenotypic changes might link to disease progression. Higher levels of MHC-I expression are known to be detrimental to β-cell survival^7^, and reduced expression levels of functional markers (e.g., TXNIP) could establish a heightened susceptibility of β-cells to stress pathways. On the other hand, lower levels of lineage markers and thus, a more de-differentiated β-cell state, might confer a survival bias but also, reduce β-cell function^55^. Although we expected signs of heightened ER stress^26,27,28^, the ER stress markers WFS-1, IRE1α, and XBP1, as well as NFκB were downregulated. We note, however, that this reduction of ER-stress and other markers could still be downstream of chronic cycles of ER stress, as this was shown to lower β-cell protein levels over time^54^. Irrespective of the mechanism, our data suggest that there is downregulation of protein levels across most markers in β-cells in T1D in addition to any marker-specific losses, which should guide future β-cell studies across disease stages.

While most β-cells in Onset donors displayed MHC-I upregulation and downregulated protein levels, β-cells within insulitic islets specifically expressed MHC-II and upregulated IFN-responsive markers. This is likely caused by a shift to an IFN-γ driven cytokine environment^7^ that we observed during the infiltration of exhausted-like T cells, which in turn may precipitate infiltration of further non-exhausted T cells, e.g. via ICAM-1 upregulation. The insulitic-specific β- cell state might facilitate direct antigen presentation between β-cells and T-CD4 cells but could also reflect β-cell protective effects^56^. While effect sizes were often largest in β-cells, we observed similar marker changes in other endocrine and islet-proximal exocrine cells both for MHC-I, with significant upregulation in α-cells already in mAAb+ donors, as well as for MHC-II. This suggests general islet-wide and islet-proximal cytokine effects both prior to and after disease onset and therefore, extends the question why β-cells, but not other endocrine or islet-proximal cells are killed^57^. Here, comparisons to α-cells, but also to δ-cells and islet-proximal exocrine cells will be informative.

In parallel, we analyzed immune cell abundances and infiltration across T1D disease stages. The earliest detected perturbations in the islet-immune interface of the human pancreas were HLA-ABC expression in β-cells, as well as infiltration of M1/M2-like act. macrophage cells and islet-proximal infiltration of T-CD4 PD1+ act. cells. These immune cell subtypes were enriched already in sAAb+ donors, and indeed were the only subtypes enriched at this early disease stage, indicating their importance for early disease progression and suggesting that they may interact. Consistent with this, in mAAb+ and Onset donors, we observed several indicators of interactions between myeloid cells (mainly M1/M2-like act. macrophages and cDCs) and exhausted-like T- CD4 and T-CD8 cells (PD1+ act. cells and T-ex^e^^7^ cells), suggesting the existence of a myeloid cell- exhausted-like T cell axis. Notable interactions were observed both at the islet-edge in mAAb+ donors and during overt islet infiltration, key stages of disease progression^58^. We also detected co-stimulatory and APC phenotypes, such as M1-polarized macrophages and mature cDCs, suggesting antigen-presentation and co-stimulation to T cells^21,59^. These signals might be required to deliver survival signals to T cells and/or subsequent effector program acquisition for precursor exhausted T cells (T-PEX), which have been attributed a key role in T1D and might be comprised within the here described exhausted-like T cell populations^60,61,62,21^. Nevertheless, while these interactions imply diabetogenic effects, the high levels of TIM-3 on cDCs suggest an immuno-regulatory phenotype that might slow T1D progression^49^, and the abundant interactions between macrophages with exhausted-like T cells also suggest a role in T cell exhaustion, as observed in tumor immunity^63,64^. Based on these data, we propose that the role of macrophages and DCs in the human pancreas to T1D extends beyond cytokine secretion and that instead, interactions between macrophages and DCs with exhausted-like T cells govern key diabetogenic but also protective effects in T1D progression.

This myeloid cell - exhausted-like T cell axis is likely targeted by immune checkpoint inhibitors in cancer therapy, which may explain the reported adverse induction of T1D^65^. This would apply not only to currently approved PD-1 treatment targeting exhausted-like T cells^66^, but also to anti-TIM- 3 treatment potentially targeting both islet-infiltrating TIM-3^+^ exhausted-like T cells and the detected TIM-3^high^ cDCs^49,67^. The T-CD4 PD1+ act. cells show lower PD-1 and especially TIM-3 levels than T-ex^e^^7^ cells, suggesting a less terminally exhausted state^68^. As substantial work has focused on auto-reactive T-CD8 cells^61,69^, it will be interesting to further decipher the role of exhausted-like T-CD4 cells, both in T1D progression^60^ and during adverse induction of T1D during checkpoint inhibitor treatment.

Younger patients present with more aggressive disease than adolescents and young adults^50^; thus, we analyzed our data for age-dependent effects. Insulitic infiltrates containing exhausted-like T cells and pro-inflammatory macrophages were present across age. This further underlines their importance to disease and highlights the clinical potential to modulate this axis in T1D patients across age to enforce exhaustion in T cells. Further, we observed that immune cells of TLS-like structures (i.e., B cells and naïve T-CD8 cells) were enriched in younger donors (<13 years) in comparison to older donors (≥ 13 years), akin to a previously proposed stratification (CD20^high^: < 7 years, CD20^low^: ≥ 13 years, mixed: 7-12 years)^45^. Nevertheless, we also detected (albeit rarely) B cell-high areas in some older donors, which could refine this stratification. While a limitation of our study is that it included fewer younger than older donors, our data are supported by previous results in which islet-associated TLS-like structures were also observed in donors older than 12 years of age^51^.

Across disease stages, including non-diabetic controls, we observed CD11c^+^ macrophages and marginally activated T-CD8s as mainly stratifying cells between younger and older donors. These differences thus likely reflect the evolution of the pancreatic immune niche composition across age, with potential implications for T1D disease. Due to high STING protein levels on CD11c+ macrophages, we hypothesize these cells to exhibit an Interferon-mediated anti-viral innate immune program in young donors, as was observed in the lung^70^. This is highly relevant (i) as viral infections modulate risk for auto-immunity onset^71^ and (ii) SNPs in innate dsRNA pattern-recognition receptors like *IFIH1* are known risk-factors for progression to T1D disease^72,74^. Further studies using healthy as well as diseased pancreata, in the context of T1D or viral infections, should establish function of these cells and verify T1D-specific effects, for example by an aberrant activation of these cells in younger donors.

Our study has limitations. First, IMC is limited by data acquisition speed, thus limiting acquisition of whole-slide field-of-views and data per tissue section. Therefore, we focused on imaging many small ROIs that included islets and proximal exocrine tissue, to strike a balance between data acquisition time and informative data content. Second, we did not derive antigen-specificity of T cells. Third, studying longitudinal progression of a disease by cross-sectional comparisons of donor tissue sections is inherently limited, since these samples are post-mortem snapshots of disease, potentially influenced by end-of-life effects and organ handling^39^. Nevertheless, it is currently the only feasible option for large-scale studies on human pancreatic tissue.

We envision that in combination with the present study, future multimodal studies of human pancreata from subjects with diverse genetic backgrounds could facilitate creation of T1D disease atlases that can be used to associate disease features with clinical co-variates and intervention outcomes and thereby help to develop personalized strategies to treat T1D.

## Resource Availability

### Lead Contact

Requests for further information and resources should be directed to and will be fulfilled by the lead contact, Bernd Bodenmiller, PhD.

### Materials Availability

Additional tissue samples from donors evaluated as a part of this study can be requested from nPOD for use in projects approved by the nPOD Tissue Prioritization Committee, as outlined on the nPOD website (https://npod.org/). This study did not generate new unique reagents.

### Data and Code Availability

We will provide the raw IMC .mcd/.txt files, .tiff image stacks, CNN islet segmentation model, cell and islet masks, as well as annotated single-cell objects at Zenodo upon publication for both pilot study and here presented original study. The code to perform IMC pre-processing, data analysis and visualization supporting this study will be available via GitHub upon publication.

## Supplemental Information Titles and Legends

Document S1. Figures S1–S10. Document S2. Tables S1-S8.

## STAR Methods

### Key Resources Table

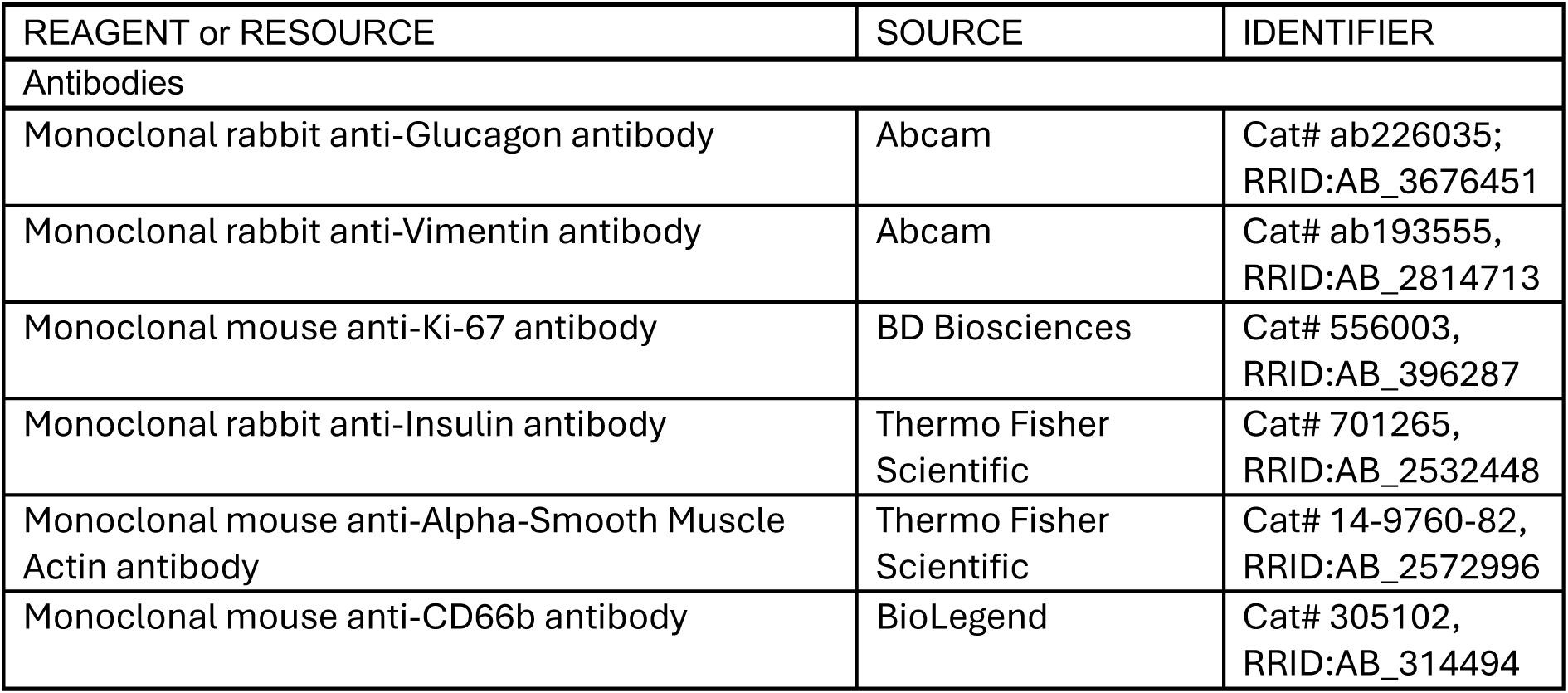
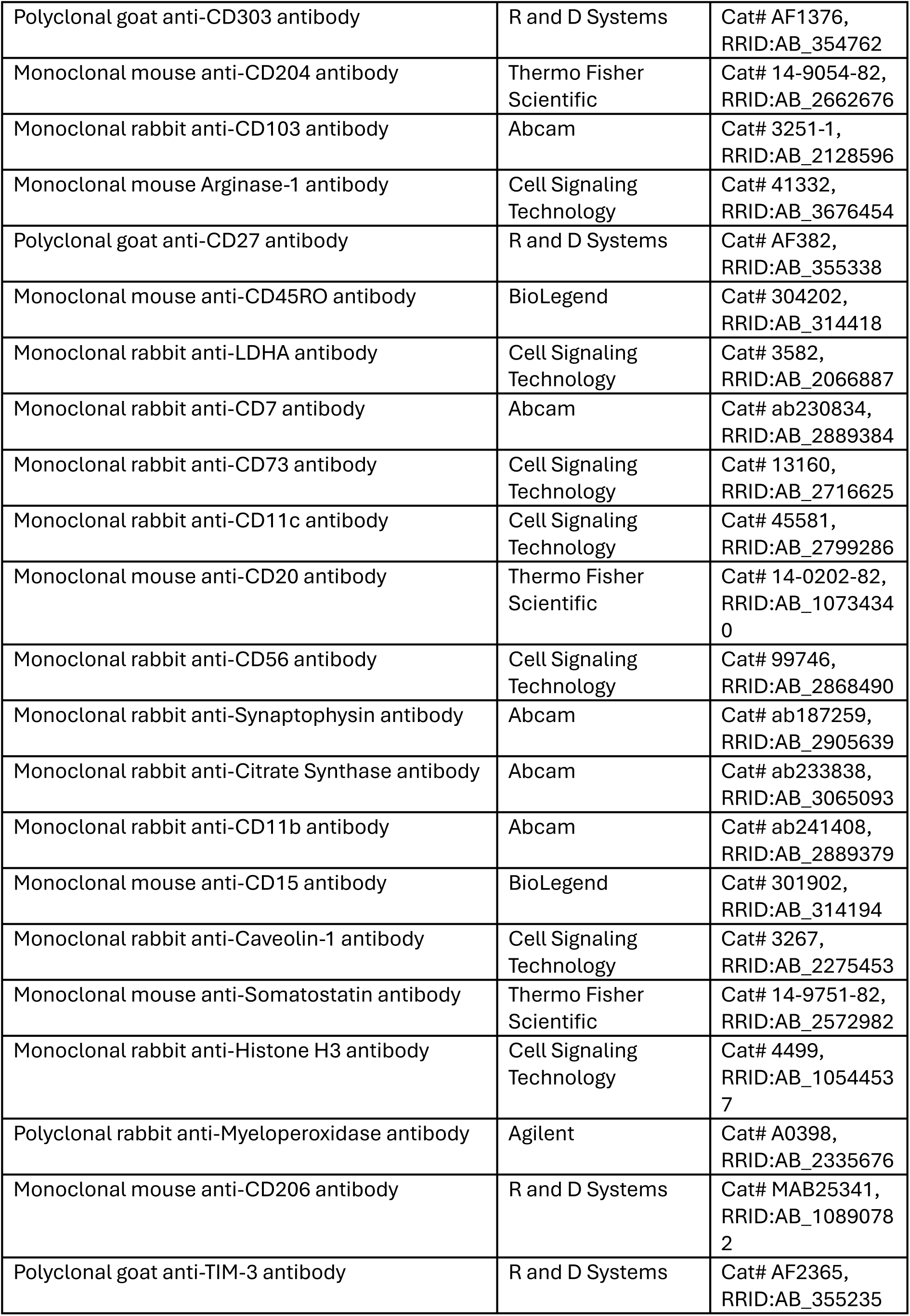
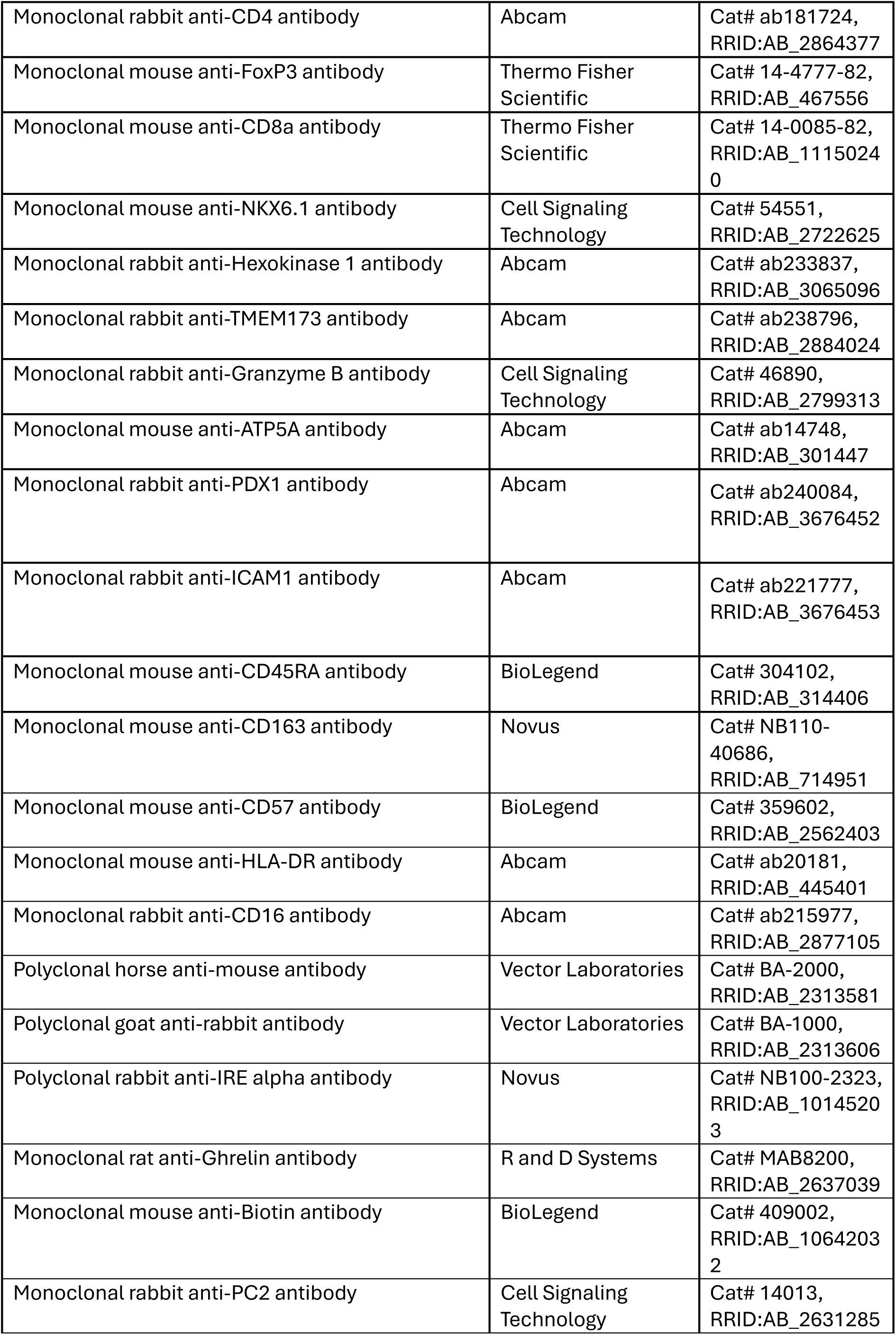
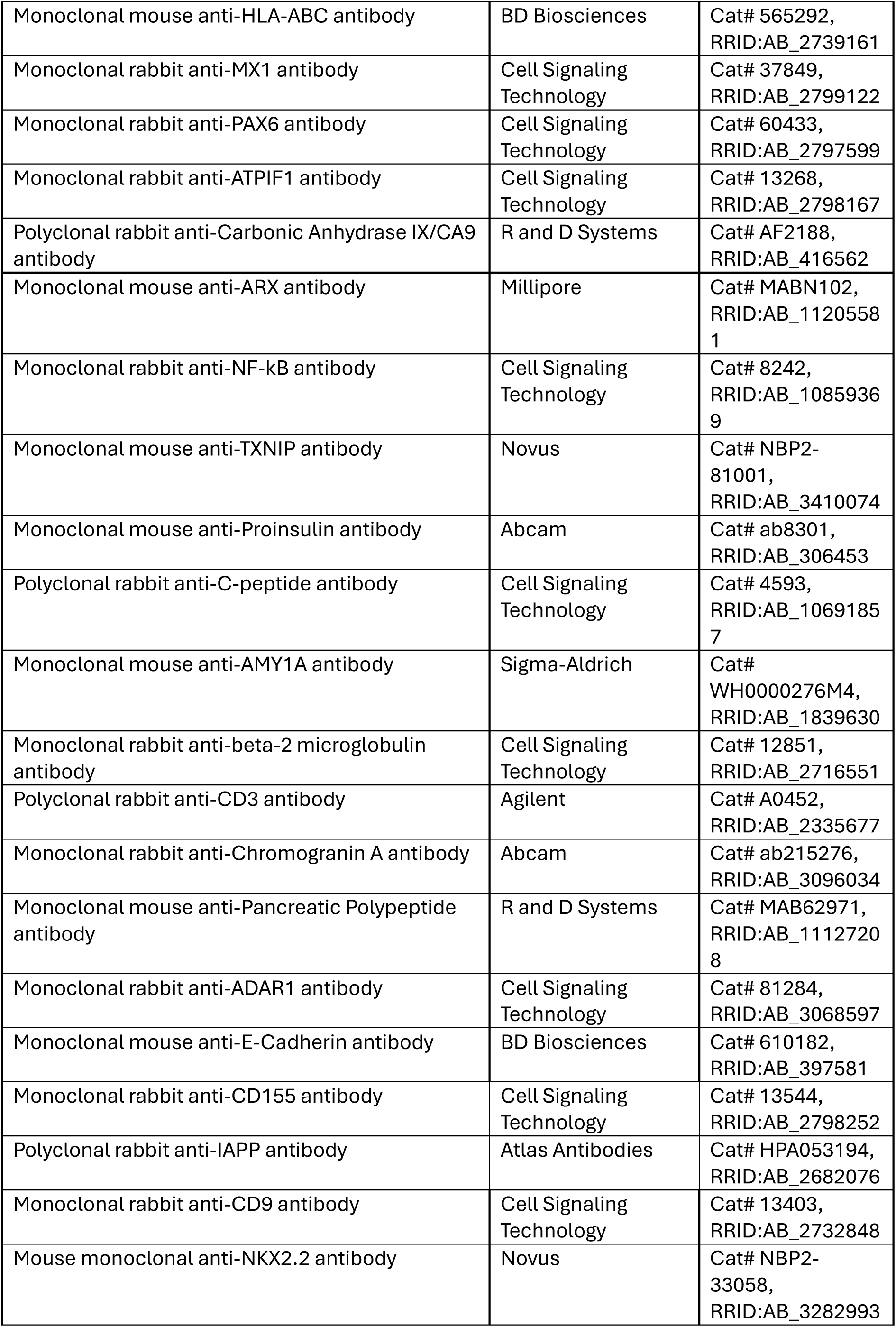
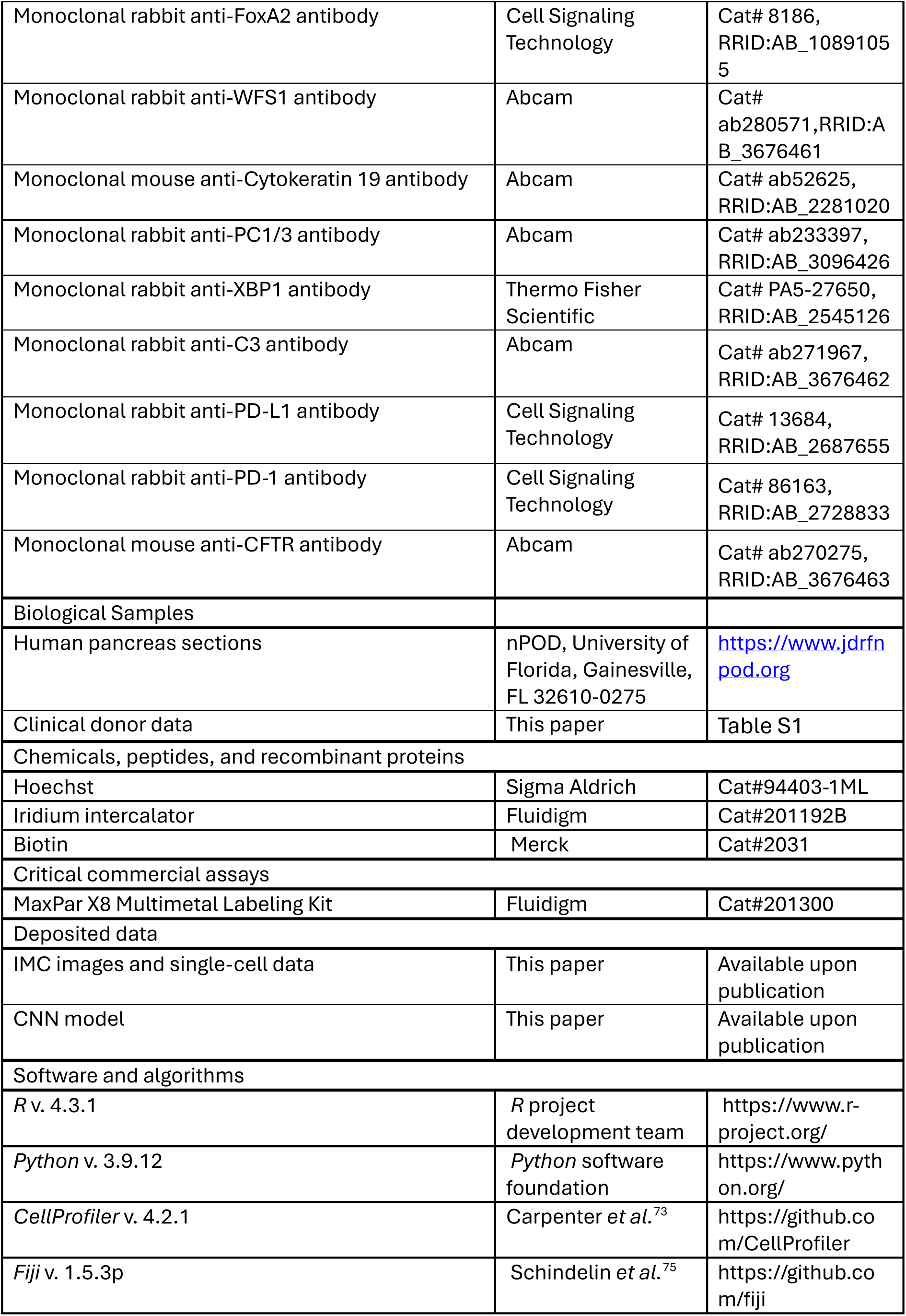
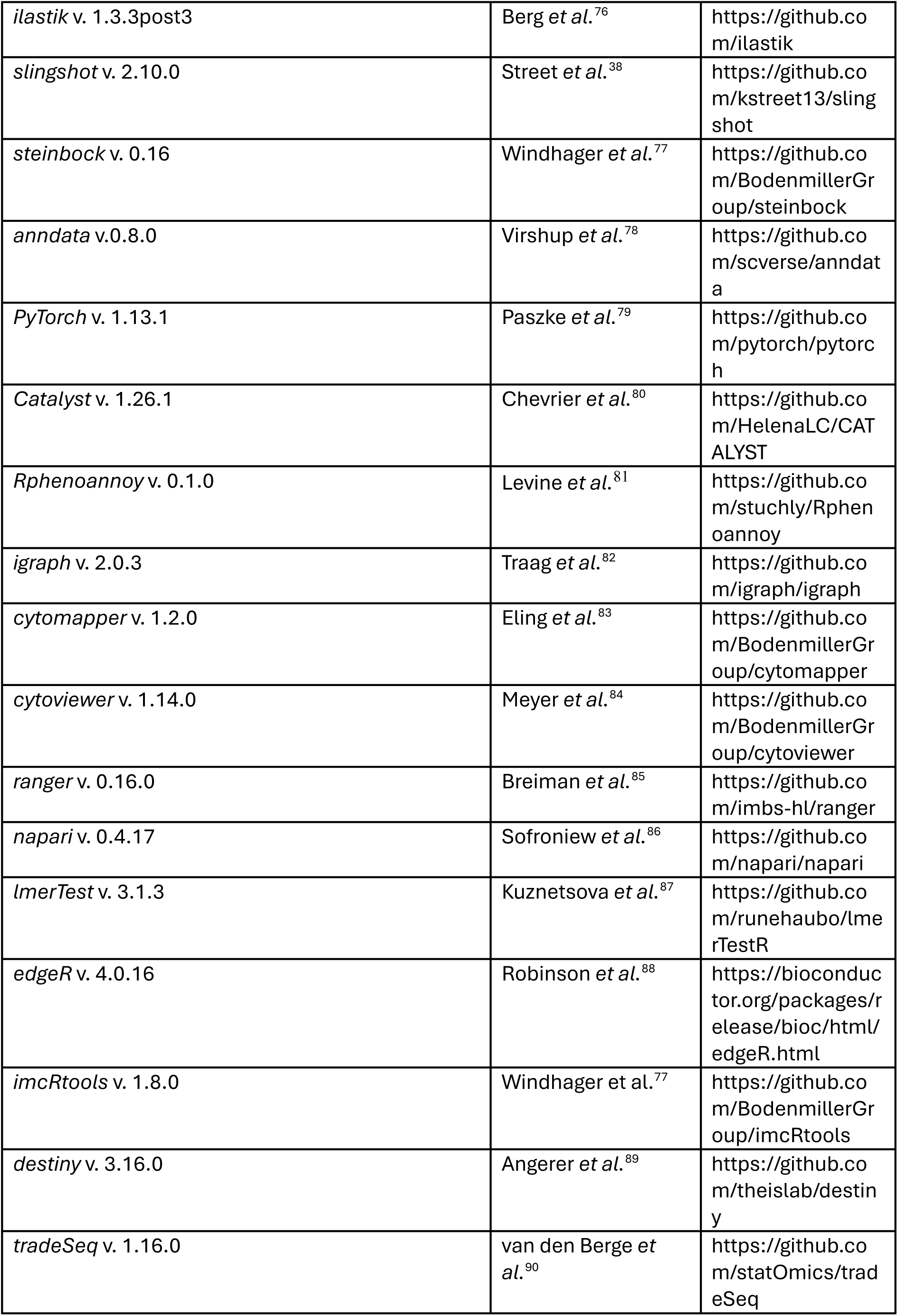
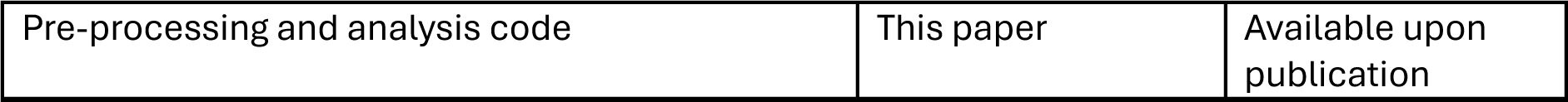

### Donor Details

Pancreatic samples were obtained from nPOD. Procedures were also approved by the United Network for Organ Sharing (UNOS) according to federal guidelines, and informed consent was obtained from each donor’s legal representative. All tissue samples, as well as corresponding clinical information were collected after approval by the University of Florida Institutional Review Board (IRB201600029). Donors included subjects with sAAb+, mAAb+, recent-onset (Onset) T1D (≤2 years), long-standing (LD) T1D (≥3 years), and autoantibody negative controls without diabetes. Cases were matched by BMI, sex, and age across batches and disease stages (Figure S1). Full donor information is listed in Table S1.

## Method Details

### Antibody testing and validation

Antibodies were initially tested with immunofluorescence. Antibodies targeting immune markers were tested on lymphoid tissues (spleen, tonsil, and lymph node), and antibodies targeting pancreas markers were tested on pancreatic tissue. Markers displaying expected expression patterns were conjugated to heavy metals and tested by IMC.

### Pilot study

Formalin fixed paraffin embedded (FFPE) pancreatic tissue sections from 16 cadaveric organ donors were stained (control, N=4; sAAb+, N=4; mAAb+, N=4; Onset, N=4). A total of 130 distinct antibodies were conjugated to metals and tested by IMC in five separate panels (Table S4-8). Antibodies were selected for the inclusion in the immune or islet panel if their staining pattern was consistent with the literature, had a high signal-to-noise ratio, and was non-redundant.

### Antibody conjugation and titration

AirLab was used to design antibody panels^91^. Metals were conjugated to all antibodies using the MaxPar X8 Multimetal Labeling Kit (Fluidigm) according to the manufacturer’s instructions. FFPE pancreatic tissue sections were stained with serial dilutions of relative concentrations of 1500, 1000, 500, 250, and 125. One section was used per dilution. Pancreatic sections were imaged by IMC, raw IMC data was processed by *steinbock*^77^ (v. 0.10.2), and processed data were imported as *anndata* object^92^ (v. 0.8.0). The signal-to-noise ratio (SNR) was calculated by defining positive and negative cells. The dilution with the highest signal-to-noise ratio was selected for each antibody.

### Tissue staining

Per organ donor, two consecutive 4µm pancreatic tissue sections were stained, one with the islet panel and one with the immune panel (Table S2, 3).

FFPE tissue sections were deparaffinized and rehydrated using a graded alcohol series. Antigen-retrieval was performed in a decloaking chamber for 30 minutes at 95° C in HIER buffer (pH 9.2), and tissues were blocked with 10% normal horse serum. Sections were first incubated for 1 hour at 4°C with unconjugated primary antibodies (Immune panel: mouse anti-CD3e, rabbit anti-PD1;

Islet panel: Biotin (HBP), mouse anti-CD45RA, mouse anti-CD45RO, rabbit anti-PD-L1). Next, sections were stained for 1 hour at room temperature with conjugated secondary antibodies (Immune panel: anti-mouse IgG-^152^Sm, anti-mouse IgG-Alexa Fluor 555, anti-rabbit IgG-^166^Er; Islet panel: anti-rabbit IgG-^160^Gd, anti-mouse IgG-Alexa Fluor 555) and counterstained with Hoechst. Pancreatic tissue sections were then incubated with metal-conjugated primary antibodies at room temperature for 3.5 hours (Islet panel: 13 antibodies; Immune panel: 17 antibodies, Table S2,3) and incubated overnight at 4°C with the remaining metal-conjugated primary antibodies and mouse anti-CD99 Allophycocyanin.

At this stage, IF images were acquired (see below) and slides were counterstained with iridium DNA intercalator for 5 minutes at room temperature before air-drying using pressurized air.

### ROI selection

Brightfield and immunofluorescent images of the pancreatic sections stained with the islet and immune panels were acquired on a slide scanner (Zeiss Axio Scan.Z1) at 2.5x and 10x magnification, respectively. The immunofluorescent images of the immune panel (CD3e, CD99, DAPI) were automatically registered to the immunofluorescent images of the islet panel (CD45, CD99, DAPI) in *FiJi*^75^ (v. 1.5.3p) using the virtual stack slices registration plugin (v. 3.0.7).

ROIs were selected based on the islet panel immunofluorescent images. First, islets were segmented by training a pixel classifier in *ilastik*^76^ (v. 1.3.3post3) and labeling pixels as either islet or exocrine and background. Pixel probability maps were then exported to *CellProfiler*^73^ (v. 4.2.1) to segment islets. Islet masks were imported into *FiJi* and loaded as a set of ROIs using rectangular bounding boxes. Bounding boxes of the islet panel and immune panel were expanded by 50 µm and 80 µm, respectively. Equal tissue areas across panels for IMC ablation were selected by first recording ROI coordinates on the islet panel and then transforming the ROI coordinates to the immune panel. ROIs focused on imaging islets, their proximal exocrine tissue, and their infiltration by immune cells. Imaged islets were sampled evenly across tissue position, size distribution, and immune cell infiltration.

### IMC data acquisition

The consecutive sections, respectively stained by the islet and immune panels, were acquired on two separate Helios time-of-flight mass cytometers (CyTOF) coupled to Hyperion Imaging Systems (Fluidigm). Laser ablation was set at 200 Hz and resolution was 1 µm. Prior to acquisition, optical images of slides were acquired using the Hyperion software, and immunofluorescent ROI coordinates were translated to IMC ROI coordinates by manually selecting landmarks and performing image registration.

Data were acquired in four batches in a randomized order. Samples were assigned to batches by using a variance minimization approach to minimize inter-batch differences^93^. Performance stability was ensured by daily calibrating the mass cytometers with a tuning slide spiked with five metal elements (Fluidigm). For each section, 75 ROIs were acquired per panel. This yielded 14,050 45-plex image stacks from 88 pancreas sections.

### Quantification and Statistical Analysis

A multi-step pre-processing pipeline was implemented, which included reading of raw IMC files, islet segmentation, cell segmentation, image registration, and feature extraction (Figure S2A).

### IMC data processing

Acquired IMC data were saved as .txt and .mcd files and converted to .tiff image files using the *preprocess* functionality of *steinbock* (v. 0.15) with a hot pixel threshold of 50. Images will be available via Zenodo upon publication.

### Islet segmentation

#### Generation of binary training masks

The IMC data from the pilot study was used as training data. Here, *ilastik* and *CellProfiler* were used to segment islets from all five consecutive sections. Only channels of informative markers were used to perform pixel classification (Table S9).

#### Training of CNN-based islet segmentation model

The islet binary masks, predicted from the IMC data of the pilot study, were used to train a deep convolutional neural network (CNN). This CNN was used for downstream inference of islet segmentation masks of IMC data acquired from the patient cohort. Here, IMC ROIs were split into training (67%) and test (33%) sets, while stratifying for donor. Expression of informative channels (Table S9) were min-max normalized and aggregated. The CNN was implemented using the *PyTorch* framework^79^ (v. 1.13.1).

#### Inference and post-processing

The trained U-Net model was applied to acquired ROIs. Recorded signal intensities of informative channels of the islet panel (SYP, CHGA) and immune panel (SYP) were normalized and aggregated. Images were resized to 160 x 160 pixels, and the trained U-Net model was used to infer islet masks. Images and predicted masks were resized to their original size and region properties of individual islets were computed. Islets smaller than 50 pixels with an eccentricity above 0.95 were not considered. Binary holes in the islets were filled.

Images and masks were visually inspected, and images with (i) bad image quality, (ii) mismatched acquired regions on consecutive sections, (iii) disrupted acquisition due to software crash or experimental handling, or (iv) islets were not segmented were not included in downstream analyses. Over 11,000 islets were segmented and labeled (Figure S2B, C). The model and inferred islet masks will be available at Zenodo upon publication.

#### Cell segmentation

Whole-cell segmentation masks were generated using the pre-trained deep learning segmentation model Mesmer from the DeepCell library^94^ as implemented in *steinbock* (v. 0.16). Informative nuclear markers (DNA1, DNA3, and Histone H3) and membrane or cytoplasmic markers (Immune panel: HLA-DR, SYP, CD45RA, CD45RO, Caveolin-1, Vimentin; Islet panel: CD3e, SYP, CD163, E-Cadherin, Vimentin, Caveolin-1, E-Cadherin) were min-max-normalized, and their intensities were aggregated. The aggregated nuclear and cytoplasmic intensities were used as input for the DeepCell model. Approximately 6.2 million cells were segmented in the islet panel and approximately 10.5 million cells were segmented in the immune panel (Figure S2B, C). Cell masks will be available at Zenodo upon publication.

### Object measurements

Features were extracted using functionalities from *steinbock*^77^. The following features were extracted: (i) average marker intensities for whole-cells and islets, (ii) measurements of islets and whole-cells such as cell area, position, and eccentricity, (iii) the distance of cells to the islet edge, and (iv) cellular neighbors. Measured objects will be available at Zenodo upon publication.

### Spillover compensation

Channel crosstalk was removed on the single-cell level by performing spillover compensation^80^. For each acquired batch, a slide was spotted with the same metals used for antibody conjugation and was measured with IMC. The generated spillover matrices were used to correct signal spillover with the *Catalyst* package^80^ (v. 1.26.1) (Figure S2D).

### Data transformation and normalization

Single-cell expression data was normalized by three approaches. (i) Counts were arcsine-transformed using a co-factor of 1. (ii) The arcsine-transformed counts were further embedded using the *fastMNN* functionality from the *batchelor* package^95^ (v. 1.18.1). (iii) Counts were scaled by min-max normalization between 0 and 1. Values higher than the 99th percentile were clipped to a value of 1 to account for outliers. If not indicated otherwise, arcsine-transformed counts (i.e., approach i) were used in the downstream analysis. Approaches ii and iii were used for cell type annotation.

### Cell type annotation

To identify single-cell phenotypes, cells were clustered separately per antibody panel using a multi-step approach. If not indicated otherwise, within each step four separate annotations were performed, by combining two unsupervised clustering approaches with two distinct data normalizations.

Unsupervised clustering was carried out by two approaches: by using the PhenoGraph algorithm, as implemented in the *Rphenoannoy* package^81^ (v. 0.1.0), and by constructing a shared-nearest neighbors (SNN) graph and detecting communities using the Leiden algorithm as implemented in the *igraph* package^82^ (v. 2.0.3). Scaled counts and fastMNN-embedded counts were used as input for the PhenoGraph and the Leiden algorithm.

Single cells were manually annotated to a cell type if the same cell type was predicted in three out of four annotations. Otherwise, cells were annotated as ‘Ambiguous’. Expected marker expression of annotated cell clusters were visualized with *cytomapper*^83^ (v. 1.2.0), *cytoviewer*^84^ (v. 1.14.0), and *napari* (v. 0.4.17)^86^.

### Islet panel annotation

Islet panel annotation was performed in three steps. First, cells within the islet mask were annotated as α-, β-, δ-, ε-, γ-, and ‘Other’ cells using lineage markers of the islet and the wider tissue context (Table S10). Second, endocrine cells outside of the islet-mask were predicted and annotated using a random forest classifier as implemented in *ranger*^85,96^ (v. 0.16.0) using the previously annotated endocrine cells as labeled training data. Finally, non-endocrine cells were annotated as ductal, acinar, endothelial, mesenchymal, T cells, macrophages, and ‘Other’ using informative lineage and functional markers (Table S10).

### Immune panel annotation

Immune panel annotation was performed in five steps. First, cells were separated into either immune or non-immune cells using all markers except the nuclear channels and the proliferation marker Ki67 (Table S11). Second, all non-immune cells were annotated as α, β, δ, ‘Islet-Other’, acinar, ductal, nerve, endothelial, smooth-muscle, and ‘Other’ using lineage markers, functional markers, and metabolic markers (Table S11). Third, immune cells were separated into major immune categories (i.e., lymphocytes, neutrophils and myeloid cells) using immune and metabolic markers (Table S11). Fourth, lymphocytes were classified into B cells, NK cells, T-CD4, T-CD8, T-DN cells, and CD303^+^/VIM^+^ cells using informative lymphocytic lineage markers, functional markers, and metabolic markers. Finally, T-CD8, T-CD4, myeloid cells, neutrophils, and NK cells were sub-clustered using specific cell-type relevant markers (Table S11).

One of the main axes of variation during clustering of protein levels of T cells was total protein level, likely reflecting activation status. Here, we observed previously described high FoxP3 protein levels in both activated T-CD4 cells and activated T-CD8 cells^97^, which was exemplified by strong correlation to CD3e protein levels (T-CD8 cells, *r*=0.78; T-CD4 cells, *r*=0.67). This prevented reliable annotation of T-regs during clustering. Nevertheless, in T-CD4 but not in T-CD8 cells, we observed FoxP3^high^CD3^mid^ cells, which we assumed to be regulatory cells, but not activated conventional T cells (FoxP3^high^CD3^high^). We thus employed a strategy equivalent to setting non-linear gates on CD3e and FoxP3 expression. Here, we assigned these cells by fitting a generalized additive model to FoxP3 and CD3e expression protein levels of T-CD8 cells, the trend exemplifying the FoxP3 upregulation in activated T cells. Next, we calculated the FoxP3 residuals and assigned T-CD4 cells with high FoxP3 expression (>1 arcsinh-scaled counts) and large FoxP3 expression residuals (greater than the 99^th^ quantile of T-CD8 FoxP3 residuals), as T- regs.

### Statistical analysis

Statistical analyses were performed using *R* (v. 4.3.1). Statistical analyses are described in the figure legends. All statistical tests were adjusted for multiple comparisons by false-discovery rate (FDR) correction^98^. As 75 IMC ROIs were acquired per organ donor, data were aggregated by ROI and linear mixed-effect models (LMM) were applied to test for statistical significance between disease stages. LMMs with random intercepts were fitted using the *lmerTest* package^87^ (v. 3.1.3) with the case ID of each donor as a random effect. Otherwise, data aggregated by donor and disease stage were compared by Wilcoxon rank-sum tests. Differential abundance tests were carried out using *edgeR*^88^(v. 4.0.16).

Recent work has suggested the existence of age-associated endotypes^29^. Our cohort included donors aged <7 years (N=11), 7-12 years (N=9), and ≥13 years (N=68). We merged donors <13 years into a single category (N=20). We fit two separate LMMs to estimate the influence of age.

The first model was just applied to Onset T1D organ donors:

Y ∼ disease_duration + age_group + (1|case_id)

The second model was applied to control, sAAb+, mAAb+, and Onset T1D donors:

Y ∼ disease_stage + age_group + (1|case_id)

Y denotes marker expression, cell type density or immune cell subtype density. Disease duration is the time since onset, and age_group denotes the two age groups defined above (< 13 years; ≥13 years). We excluded LD donors from this analysis, as there was significant a time-lag between age of onset and age of donation (up to 21 years), likely skewing age-associated effects.

### Pseudotime analysis

Pseudotime analysis of β-cell expression profiles was performed using *slingshot*^38^ (v. 2.10.0). We used default parameters to identify the global lineage structure, and fit smooth branching curves to infer the pseudotime variables for each β-cell. Notably, we did not provide start and leaf nodes or cluster information and thus, *slingshot* inferred the global trajectory in an unbiased manner. As input, we used diffusion map embedded single-cell β-cell expression profiles, which were computed using *destiny*^89^ (v. 3.16.0). We considered the expression of 28 β-cell lineage and functional markers for computation of the diffusion map and pseudotime (Table S10). The association of β-cell markers to pseudotime was computed using *tradeSeq*^90^ (v. 1.16.0). We computed cross-correlation of marker expression, cell type density, and infiltration scores along pseudotime using the *ccf*-functionality of the *stats* package. We computed Granger causality using the *grangertest*-functionality of *lmtest (v. 0.9.40)* to estimate sequencing of events along pseudotime.

### Infiltration score

The infiltration score was assigned to every cell by assigning a value of ‘1’ to islet-infiltrating cells and a value of ‘1/d’ to cells outside the islet. Here, *d* is the distance from the islet edge in µm. The infiltration score was further normalized by the total number of cells per image. By weighting for distance, this score rewards abundance increases in the peri-islet space and penalizes abundance increases in tissue areas distal to the islet.

### Spatial graph

Two spatial neighborhood graphs were built for each image by using *imcRtools*^77^ (v. 1.8.0). One contained the 5 nearest neighbors within a 15-µm radius, and one graph considered the 10 nearest neighbors within a 25-µm radius. Cells 10 µm from the image edge were removed to exclude edge effects. *imcRtools* was used both to aggregate the cellular composition and expression of neighboring cells.

### Cellular neighborhood analysis

We applied CNs analysis to the immune panel data to identify spatial tissue areas of defined cell type composition^47^. We used *k*-means clustering to aggregate the cellular composition of the 10 nearest-neighbors of each cell. Given that highly relevant biological processes such as T cell infiltration are rare and likely change across disease progression, we hypothesized that a high number of CNs would be required to capture the biological complexity of T1D progression. Therefore, we tested multiple settings for *k*. We estimated cluster stability with the Silhouette score and tested for biological relevance by calculating the association of each CN fraction to disease progression and identified *k*=60 as the ‘optimal’ setting. CNs were manually annotated by their enriched cell types and their localization relative to the islet. We merged CNs if we observed homogeneous cell type distributions between clusters, and similar fractions across disease stages. We explicitly did not merge T cell-enriched CNs, as we assumed their relevance for disease.

## Acknowledgements

We thank the organ donors and their families. This study was supported by the following grants from the National Institutes of Health (NIH): R01DK131059, R01DK123292, and P01AI042288 to MAA, and U01DK135001 to CHW. This research was performed with the support of the Network for Pancreatic Organ donors with Diabetes (nPOD; RRID:SCR_014641), a collaborative type 1 diabetes research project supported by Breakthrough T1D and The Leona M. & Harry B. Helmsley Charitable Trust (Grant# 3-SRA-2023-1417-S-B). The content and views expressed are the responsibility of the authors and do not necessarily reflect the official view of nPOD. Organ Procurement Organizations (OPO) partnering with nPOD to provide research resources are listed at https://npod.org/for-partners/npod-partners/. DMD is supported by a postdoctoral fellowship from Breakthrough-T1D (3-PDF-2025-1675-A-N). We thank the University of Zurich, for access to virtual machines and high-performance computing clusters. BB acknowledges the support of the EFSD/JDRF/Lilly Programme on Type 1 Diabetes Research. BB was funded by three SNSF grants (SNF project grant 310030_205007, SNF R’Equip grant 316030_213512, Innosuisse Bridge Grant 40B2-0_203478), an SPHN/PHRT Swiss Precision Oncology National Data Stream grant, Tumor Profiler Center funding, Skintegrity funding, Comprehensive Cancer Center Zürich Precision Oncology project funding, an NIH grant (UC4DK108132), the European Research Council (ERC) under the European Union’s Horizon 2020 Program under the ERC grant agreement no. 866074 (“Precision Motifs”). Graphical abstracts depicting the workflow were created with BioRender (biorender.com).

## Author contributions

Conceptualization: M.A.A., B.B.; Supervision: M.A.A., B.B.; Resources: I.K., M.A.A., B.B.; Funding acquisition: C.H.W., M.A.A., B.B.; Data Curation: N.S., N.D.; Formal Analysis: N.S., N.D.; Investigation: N.S., N.D., S.E.; Methodology: N.S., N.D.; Project Administration: N.S., N.D.; Writing: Original draft N.S.; Writing: Review & Editing: D.M.D., M.D.W., A.L.P., M.A.B., T.M.B., C.H.W., M.A.A., N.D.S., B.B.

## References

1. DiMeglio, L.A., Evans-Molina, C., and Oram, R.A. (2018). Type 1 diabetes. The Lancet 391, 2449–2462. 10.1016/S0140-6736(18)31320-5.

2. Ong, K.L., Stafford, L.K., McLaughlin, S.A., Boyko, E.J., Vollset, S.E., Smith, A.E., Dalton, B.E., Duprey, J., Cruz, J.A., Hagins, H., et al. (2023). Global, regional, and national burden of diabetes from 1990 to 2021, with projections of prevalence to 2050: a systematic analysis for the Global Burden of Disease Study 2021. The Lancet 402, 203–234. 10.1016/S0140-6736(23)01301-6.

3. Atkinson, M.A., and Mirmira, R.G. (2023). The pathogenic “symphony” in type 1 diabetes: A disorder of the immune system, β cells, and exocrine pancreas. Cell Metabolism 35, 1500– 1518. 10.1016/j.cmet.2023.06.018.

4. Ziegler, A.G., Rewers, M., Simell, O., Simell, T., Lempainen, J., Steck, A., Winkler, C., Ilonen, J., Veijola, R., Knip, M., et al. (2013). Seroconversion to Multiple Islet Autoantibodies and Risk of Progression to Diabetes in Children. JAMA 309, 2473–2479. 10.1001/jama.2013.6285.

5. Insel, R.A., Dunne, J.L., Atkinson, M.A., Chiang, J.L., Dabelea, D., Gottlieb, P.A., Greenbaum, C.J., Herold, K.C., Krischer, J.P., Lernmark, Å., et al. (2015). Staging Presymptomatic Type 1 Diabetes: A Scientific Statement of JDRF, the Endocrine Society, and the American Diabetes Association. Diabetes Care 38, 1964–1974. 10.2337/dc15-1419.

6. Damond, N., Engler, S., Zanotelli, V.R.T., Schapiro, D., Wasserfall, C.H., Kusmartseva, I., Nick, H.S., Thorel, F., Herrera, P.L., Atkinson, M.A., et al. (2019). A Map of Human Type 1 Diabetes Progression by Imaging Mass Cytometry. Cell Metab 29, 755–768.e5. 10.1016/j.cmet.2018.11.014.

7. Herold, K.C., Delong, T., Perdigoto, A.L., Biru, N., Brusko, T.M., and Walker, L.S.K. (2024). The immunology of type 1 diabetes. Nat Rev Immunol 24, 435–451. 10.1038/s41577-023-00985-4.

8. Coppieters, K.T., Dotta, F., Amirian, N., Campbell, P.D., Kay, T.W.H., Atkinson, M.A., Roep, B.O., and von Herrath, M.G. (2012). Demonstration of islet-autoreactive CD8 T cells in insulitic lesions from recent onset and long-term type 1 diabetes patients. J Exp Med 209, 51–60. 10.1084/jem.20111187.

9. Babon, J.A.B., DeNicola, M.E., Blodgett, D.M., Crèvecoeur, I., Buttrick, T.S., Maehr, R., Bottino, R., Naji, A., Kaddis, J., Elyaman, W., et al. (2016). Analysis of self-antigen specificity of islet-infiltrating T cells from human donors with type 1 diabetes. Nat Med 22, 1482–1487. 10.1038/nm.4203.

10. Haller, M.J., Schatz, D.A., Skyler, J.S., Krischer, J.P., Bundy, B.N., Miller, J.L., Atkinson, M.A., Becker, D.J., Baidal, D., DiMeglio, L.A., et al. (2018). Low-Dose Anti-Thymocyte Globulin (ATG) Preserves β-Cell Function and Improves HbA1c in New-Onset Type 1 Diabetes. Diabetes Care 41, 1917–1925. 10.2337/dc18-0494.

11. Herold, K.C., Bundy, B.N., Long, S.A., Bluestone, J.A., DiMeglio, L.A., Dufort, M.J., Gitelman, S.E., Gottlieb, P.A., Krischer, J.P., Linsley, P.S., et al. (2019). An Anti-CD3 Antibody, Teplizumab, in Relatives at Risk for Type 1 Diabetes. N Engl J Med 381, 603–613. 10.1056/NEJMoa1902226.

12. Herold, K.C., Hagopian, W., Auger, J.A., Poumian-Ruiz, E., Taylor, L., Donaldson, D., Gitelman, S.E., Harlan, D.M., Xu, D., Zivin, R.A., et al. (2002). Anti-CD3 monoclonal antibody in new-onset type 1 diabetes mellitus. N Engl J Med 346, 1692–1698. 10.1056/NEJMoa012864.

13. Chatenoud, L., Thervet, E., Primo, J., and Bach, J.F. (1994). Anti-CD3 antibody induces long-term remission of overt autoimmunity in nonobese diabetic mice. Proc Natl Acad Sci U S A 91, 123–127.

14. Rigby, M.R., Harris, K.M., Pinckney, A., DiMeglio, L.A., Rendell, M.S., Felner, E.I., Dostou, J.M., Gitelman, S.E., Griffin, K.J., Tsalikian, E., et al. (2015). Alefacept provides sustained clinical and immunological effects in new-onset type 1 diabetes patients. J Clin Invest 125, 3285–3296. 10.1172/JCI81722.

15. Orban, T., Bundy, B., Becker, D.J., DiMeglio, L.A., Gitelman, S.E., Goland, R., Gottlieb, P.A., Greenbaum, C.J., Marks, J.B., Monzavi, R., et al. (2011). Co-Stimulation Modulation with Abatacept in Patients with Recent-Onset Type 1 Diabetes: A Randomised Double-Masked Controlled Trial. Lancet 378, 412–419. 10.1016/S0140-6736(11)60886-6.

16. Carrero, J.A., McCarthy, D.P., Ferris, S.T., Wan, X., Hu, H., Zinselmeyer, B.H., Vomund, A.N., and Unanue, E.R. (2017). Resident macrophages of pancreatic islets have a seminal role in the initiation of autoimmune diabetes of NOD mice. Proc Natl Acad Sci U S A 114, E10418– E10427. 10.1073/pnas.1713543114.

17. Ferris, S.T., Carrero, J.A., Mohan, J.F., Calderon, B., Murphy, K.M., and Unanue, E.R. (2014). A Minor Subset of *Batf3*-Dependent Antigen-Presenting Cells in Islets of Langerhans Is Essential for the Development of Autoimmune Diabetes. Immunity 41, 657–669. 10.1016/j.immuni.2014.09.012.

18. Hussain, M.J., Peakman, M., Gallati, H., Lo, S.S., Hawa, M., Viberti, G.C., Watkins, P.J., Leslie, R.D., and Vergani, D. (1996). Elevated serum levels of macrophage-derived cytokines precede and accompany the onset of IDDM. Diabetologia 39, 60–69. 10.1007/BF00400414.

19. Friedman, R.S., Lindsay, R.S., Lilly, J.K., Nguyen, V., Sorensen, C.M., Jacobelli, J., and Krummel, M.F. (2014). An evolving autoimmune microenvironment regulates the quality of effector T cell restimulation and function. Proc Natl Acad Sci U S A 111, 9223–9228. 10.1073/pnas.1322193111.

20. Lindsay, R.S., Corbin, K., Mahne, A., Levitt, B.E., Gebert, M.J., Wigton, E.J., Bradley, B.J., Haskins, K., Jacobelli, J., Tang, Q., et al. (2015). Antigen recognition in the islets changes with progression of autoimmune islet infiltration. J Immunol 194, 522–530. 10.4049/jimmunol.1400626.

21. Zakharov, P.N., Hu, H., Wan, X., and Unanue, E.R. (2020). Single-cell RNA sequencing of murine islets shows high cellular complexity at all stages of autoimmune diabetes. J Exp Med 217, e20192362. 10.1084/jem.20192362.

22. Zirpel, H., and Roep, B.O. (2021). Islet-Resident Dendritic Cells and Macrophages in Type 1 Diabetes: In Search of Bigfoot’s Print. Front. Endocrinol. 12. 10.3389/fendo.2021.666795.

23. Bottazzo, G.F., Dean, B.M., McNally, J.M., MacKay, E.H., Swift, P.G.F., and Gamble, D.R. (1985). In Situ Characterization of Autoimmune Phenomena and Expression of HLA Molecules in the Pancreas in Diabetic Insulitis. New England Journal of Medicine 313, 353–360. 10.1056/NEJM198508083130604.

24. Foulis, A.K., Farquharson, M.A., and Meager, A. (1987). Immunoreactive α-Interferon in Insulin-secreting β cells in type 1 diabetes mellitus. The Lancet 330, 1423–1427. 10.1016/S0140-6736(87)91128-7.

25. James, E.A., Joglekar, A.V., Linnemann, A.K., Russ, H.A., and Kent, S.C. (2023). The beta cell-immune cell interface in type 1 diabetes (T1D). Mol Metab 78, 101809. 10.1016/j.molmet.2023.101809.

26. Marroqui, L., Dos Santos, R.S., Op de Beeck, A., Coomans de Brachène, A., Marselli, L., Marchetti, P., and Eizirik, D.L. (2017). Interferon-α mediates human beta cell HLA class I overexpression, endoplasmic reticulum stress and apoptosis, three hallmarks of early human type 1 diabetes. Diabetologia 60, 656–667. 10.1007/s00125-016-4201-3.

27. Marhfour, I., Lopez, X.M., Lefkaditis, D., Salmon, I., Allagnat, F., Richardson, S.J., Morgan, N.G., and Eizirik, D.L. (2012). Expression of endoplasmic reticulum stress markers in the islets of patients with type 1 diabetes. Diabetologia 55, 2417–2420. 10.1007/s00125-012-2604-3.

28. Tersey, S.A., Nishiki, Y., Templin, A.T., Cabrera, S.M., Stull, N.D., Colvin, S.C., Evans-Molina, C., Rickus, J.L., Maier, B., and Mirmira, R.G. (2012). Islet β-cell endoplasmic reticulum stress precedes the onset of type 1 diabetes in the nonobese diabetic mouse model. Diabetes 61, 818–827. 10.2337/db11-1293.

29. Battaglia, M., Ahmed, S., Anderson, M.S., Atkinson, M.A., Becker, D., Bingley, P.J., Bosi, E., Brusko, T.M., DiMeglio, L.A., Evans-Molina, C., et al. (2020). Introducing the Endotype Concept to Address the Challenge of Disease Heterogeneity in Type 1 Diabetes. Diabetes Care 43, 5–12. 10.2337/dc19-0880.

30. Fasolino, M., Schwartz, G.W., Patil, A.R., Mongia, A., Golson, M.L., Wang, Y.J., Morgan, A., Liu, C., Schug, J., Liu, J., et al. (2022). Single-cell multi-omics analysis of human pancreatic islets reveals novel cellular states in type 1 diabetes. Nat Metab 4, 284–299. 10.1038/s42255-022-00531-x.

31. Wang, Y.J., Traum, D., Schug, J., Gao, L., Liu, C., Atkinson, M.A., Powers, A.C., Feldman, M.D., Naji, A., Chang, K.-M., et al. (2019). Multiplexed In Situ Imaging Mass Cytometry Analysis of the Human Endocrine Pancreas and Immune System in Type 1 Diabetes. Cell Metabolism 29, 769–783.e4. 10.1016/j.cmet.2019.01.003.

32. Barlow, G.L., Schürch, C.M., Bhate, S.S., Phillips, D., Young, A., Dong, S., Martinez, H., Kaber, G., Nagy, N., Ramachandran, S., et al. (2024). The Extra-Islet Pancreas Supports Autoimmunity in Human Type 1 Diabetes. Preprint at medRxiv, 10.1101/2023.03.15.23287145

33. K, M., F, O., F, A., and Ak, C. (2016). Endoplasmic reticulum stress and the unfolded protein response in pancreatic islet inflammation. Journal of molecular endocrinology 57. 10.1530/JME-15-0306.

34. Ml, C. F. S., and Dl, E. (2020). Molecular Footprints of the Immune Assault on Pancreatic Beta Cells in Type 1 Diabetes. Frontiers in endocrinology 11. 10.3389/fendo.2020.568446.

35. Liu, L., You, X., Han, S., Sun, Y., Zhang, J., and Zhang, Y. (2021). CD155/TIGIT, a novel immune checkpoint in human cancers (Review). Oncol Rep 45, 835–845. 10.3892/or.2021.7943.

36. Choi, E.-H., and Park, S.-J. (2023). TXNIP: A key protein in the cellular stress response pathway and a potential therapeutic target. Exp Mol Med 55, 1348–1356. 10.1038/s12276-023-01019-8.

37. Forlenza, G.P., McVean, J., Beck, R.W., Bauza, C., Bailey, R., Buckingham, B., DiMeglio, L.A., Sherr, J.L., Clements, M., Neyman, A., et al. (2023). Effect of Verapamil on Pancreatic Beta Cell Function in Newly Diagnosed Pediatric Type 1 Diabetes: A Randomized Clinical Trial. JAMA 329, 990–999. 10.1001/jama.2023.2064.

38. Street, K., Risso, D., Fletcher, R.B., Das, D., Ngai, J., Yosef, N., Purdom, E., and Dudoit, S. (2018). Slingshot: cell lineage and pseudotime inference for single-cell transcriptomics. BMC Genomics 19, 477. 10.1186/s12864-018-4772-0.

39. Campbell-Thompson, M. (2015). Organ donor specimens: What can they tell us about type 1 diabetes? Pediatr Diabetes 16, 320–330. 10.1111/pedi.12286.

40. Stridsberg, M., Sandler, S., and Wilander, E. (1993). Cosecretion of islet amyloid polypeptide (IAPP) and insulin from isolated rat pancreatic islets following stimulation or inhibition of beta-cell function. Regul Pept 45, 363–370. 10.1016/0167-0115(93)90362-c.

41. Foulis, A.K., Farquharson, M.A., and Hardman, R. (1987). Aberrant expression of Class II major histocompatibility complex molecules by B cells and hyperexpression of Class I major histocompatibility complex molecules by insulin containing islets in Type 1 (insulin-dependent) diabetes mellitus. Diabetologia 30, 333–343. 10.1007/BF00299027.

42. Quesada-Masachs, E., Zilberman, S., Rajendran, S., Chu, T., McArdle, S., Kiosses, W.B., Lee, J.-H.M., Yesildag, B., Benkahla, M.A., Pawlowska, A., et al. (2022). Upregulation of HLA class II in pancreatic beta cells from organ donors with type 1 diabetes. Diabetologia 65, 387–401. 10.1007/s00125-021-05619-9.

43. Russell, M.A., Redick, S.D., Blodgett, D.M., Richardson, S.J., Leete, P., Krogvold, L., Dahl-Jørgensen, K., Bottino, R., Brissova, M., Spaeth, J.M., et al. (2019). HLA Class II Antigen Processing and Presentation Pathway Components Demonstrated by Transcriptome and Protein Analyses of Islet β-Cells From Donors With Type 1 Diabetes. Diabetes 68, 988–1001. 10.2337/db18-0686.

44. Martin, T.M., Burke, S.J., Wasserfall, C.H., and Collier, J.J. (2023). Islet beta-cells and intercellular adhesion molecule-1 (ICAM-1): Integrating immune responses that influence autoimmunity and graft rejection. Autoimmunity Reviews 22, 103414. 10.1016/j.autrev.2023.103414.

45. Leete, P., Willcox, A., Krogvold, L., Dahl-Jørgensen, K., Foulis, A.K., Richardson, S.J., and Morgan, N.G. (2016). Differential Insulitic Profiles Determine the Extent of β-Cell Destruction and the Age at Onset of Type 1 Diabetes. Diabetes 65, 1362–1369. 10.2337/db15-1615.

46. Tietscher, S., Wagner, J., Anzeneder, T., Langwieder, C., Rees, M., Sobottka, B., de Souza, N., and Bodenmiller, B. (2023). A comprehensive single-cell map of T cell exhaustion-associated immune environments in human breast cancer. Nat Commun 14, 98. 10.1038/s41467-022-35238-w.

47. Schürch, C.M., Bhate, S.S., Barlow, G.L., Phillips, D.J., Noti, L., Zlobec, I., Chu, P., Black, S., Demeter, J., McIlwain, D.R., et al. (2020). Coordinated Cellular Neighborhoods Orchestrate Antitumoral Immunity at the Colorectal Cancer Invasive Front. Cell 182, 1341–1359.e19. 10.1016/j.cell.2020.07.005.

48. Hasegawa, H., and Matsumoto, T. (2018). Mechanisms of Tolerance Induction by Dendritic Cells In Vivo. Front Immunol 9, 350. 10.3389/fimmu.2018.00350.

49. Dixon, K.O., Tabaka, M., Schramm, M.A., Xiao, S., Tang, R., Dionne, D., Anderson, A.C., Rozenblatt-Rosen, O., Regev, A., and Kuchroo, V.K. (2021). TIM-3 restrains anti-tumour immunity by regulating inflammasome activation. Nature 595, 101–106. 10.1038/s41586-021-03626-9.

50. Redondo, M.J., and Morgan, N.G. (2023). Heterogeneity and endotypes in type 1 diabetes mellitus. Nat Rev Endocrinol 19, 542–554. 10.1038/s41574-023-00853-0.

51. Korpos, É., Kadri, N., Loismann, S., Findeisen, C.R., Arfuso, F., Burke, G.W., Richardson, S.J., Morgan, N.G., Bogdani, M., Pugliese, A., et al. (2021). Identification and characterisation of tertiary lymphoid organs in human type 1 diabetes. Diabetologia 64, 1626–1641. 10.1007/s00125-021-05453-z.

52. Vatner, R.E., and Janssen, E.M. (2019). STING, DCs and the link between innate and adaptive tumor immunity. Mol Immunol 110, 13–23. 10.1016/j.molimm.2017.12.001.

53. Cao, P., Abedini, A., and Raleigh, D.P. (2013). Aggregation of islet amyloid polypeptide: from physical chemistry to cell biology. Curr Opin Struct Biol 23, 82–89. 10.1016/j.sbi.2012.11.003.

54. Chen, C.-W., Guan, B.-J., Alzahrani, M.R., Gao, Z., Gao, L., Bracey, S., Wu, J., Mbow, C.A., Jobava, R., Haataja, L., et al. (2022). Adaptation to chronic ER stress enforces pancreatic β- cell plasticity. Nat Commun 13, 4621. 10.1038/s41467-022-32425-7.

55. Rui, J., Deng, S., Arazi, A., Perdigoto, A.L., Liu, Z., and Herold, K.C. (2017). β Cells that Resist Immunological Attack Develop during Progression of Autoimmune Diabetes in NOD Mice. Cell Metab 25, 727–738. 10.1016/j.cmet.2017.01.005.

56. Colli, M.L., Hill, J.L.E., Marroquí, L., Chaffey, J., Dos Santos, R.S., Leete, P., Coomans de Brachène, A., Paula, F.M.M., Op de Beeck, A., Castela, A., et al. (2018). PDL1 is expressed in the islets of people with type 1 diabetes and is up-regulated by interferons-α and-γ via IRF1 induction. EBioMedicine 36, 367–375. 10.1016/j.ebiom.2018.09.040.

57. Eizirik, D.L., Szymczak, F., and Mallone, R. (2023). Why does the immune system destroy pancreatic β-cells but not α-cells in type 1 diabetes? Nat Rev Endocrinol 19, 425–434. 10.1038/s41574-023-00826-3.

58. Korpos, É., Kadri, N., Kappelhoff, R., Wegner, J., Overall, C.M., Weber, E., Holmberg, D., Cardell, S., and Sorokin, L. (2013). The Peri-islet Basement Membrane, a Barrier to Infiltrating Leukocytes in Type 1 Diabetes in Mouse and Human. Diabetes 62, 531–542. 10.2337/db12-0432.

59. Unanue, E.R., Ferris, S.T., and Carrero, J.A. (2016). The role of islet antigen presenting cells and the presentation of insulin in the initiation of autoimmune diabetes in the NOD mouse. Immunol Rev 272, 183–201. 10.1111/imr.12430.

60. Aljobaily, N., Allard, D., Perkins, B., Raugh, A., Galland, T., Jing, Y., Stephens, W.Z., Bettini, M.L., Hale, J.S., and Bettini, M. (2024). Autoimmune CD4+ T cells fine-tune TCF1 expression to maintain function and survive persistent antigen exposure during diabetes. Immunity 57, 2583–2596.e6. 10.1016/j.immuni.2024.09.016.

61. Gearty, S.V., Dündar, F., Zumbo, P., Espinosa-Carrasco, G., Shakiba, M., Sanchez-Rivera, F.J., Socci, N.D., Trivedi, P., Lowe, S.W., Lauer, P., et al. (2022). An autoimmune stem-like CD8 T cell population drives type 1 diabetes. Nature 602, 156–161. 10.1038/s41586-021-04248-x.

62. Prokhnevska, N., Cardenas, M.A., Valanparambil, R.M., Sobierajska, E., Barwick, B.G., Jansen, C., Reyes Moon, A., Gregorova, P., delBalzo, L., Greenwald, R., et al. (2023). CD8+ T cell activation in cancer comprises an initial activation phase in lymph nodes followed by effector differentiation within the tumor. Immunity 56, 107–124.e5. 10.1016/j.immuni.2022.12.002.

63. Kersten, K., Hu, K.H., Combes, A.J., Samad, B., Harwin, T., Ray, A., Rao, A.A., Cai, E., Marchuk, K., Artichoker, J., et al. (2022). Spatiotemporal co-dependency between macrophages and exhausted CD8+ T cells in cancer. Cancer Cell 40, 624–638.e9. 10.1016/j.ccell.2022.05.004.

64. Nixon, B.G., Kuo, F., Ji, L., Liu, M., Capistrano, K., Do, M., Franklin, R.A., Wu, X., Kansler, E.R., Srivastava, R.M., et al. (2022). Tumor-associated macrophages expressing the transcription factor IRF8 promote T cell exhaustion in cancer. Immunity 55, 2044–2058.e5. 10.1016/j.immuni.2022.10.002.

65. Sznol, M., Postow, M.A., Davies, M.J., Pavlick, A.C., Plimack, E.R., Shaheen, M., Veloski, C., and Robert, C. (2017). Endocrine-related adverse events associated with immune checkpoint blockade and expert insights on their management. Cancer Treatment Reviews 58, 70–76. 10.1016/j.ctrv.2017.06.002.

66. Hu, H., Zakharov, P.N., Peterson, O.J., and Unanue, E.R. (2020). Cytocidal macrophages in symbiosis with CD4 and CD8 T cells cause acute diabetes following checkpoint blockade of PD-1 in NOD mice. Proc. Natl. Acad. Sci. U.S.A. 117, 31319–31330. 10.1073/pnas.2019743117.

67. de Mingo Pulido, Á., Gardner, A., Hiebler, S., Soliman, H., Rugo, H.S., Krummel, M.F., Coussens, L.M., and Ruffell, B. (2018). TIM-3 regulates CD103+ dendritic cell function and response to chemotherapy in breast cancer. Cancer Cell 33, 60–74.e6. 10.1016/j.ccell.2017.11.019.

68. Gebhardt, T., Park, S.L., and Parish, I.A. (2023). Stem-like exhausted and memory CD8+ T cells in cancer. Nat Rev Cancer 23, 780–798. 10.1038/s41568-023-00615-0.

69. Abdelsamed, H.A., Zebley, C.C., Nguyen, H., Rutishauser, R.L., Fan, Y., Ghoneim, H.E., Crawford, J.C., Alfei, F., Alli, S., Ribeiro, S.P., et al. (2020). Beta cell-specific CD8+ T cells maintain stem cell memory-associated epigenetic programs during type 1 diabetes. Nat Immunol 21, 578–587. 10.1038/s41590-020-0633-5.

70. Loske, J., Röhmel, J., Lukassen, S., Stricker, S., Magalhães, V.G., Liebig, J., Chua, R.L., Thürmann, L., Messingschlager, M., Seegebarth, A., et al. (2022). Pre-activated antiviral innate immunity in the upper airways controls early SARS-CoV-2 infection in children. Nat Biotechnol 40, 319–324. 10.1038/s41587-021-01037-9.

71. The Environmental Determinants of Diabetes in the Young (TEDDY) Study (2008). Ann N Y Acad Sci 1150, 1–13. 10.1196/annals.1447.062.

72. Steck, A.K., Dong, F., Wong, R., Fouts, A., Liu, E., Romanos, J., Wijmenga, C., Norris, J.M., and Rewers, M.J. (2014). Improving prediction of type 1 diabetes by testing non-HLA genetic variants in addition to HLA markers. Pediatr Diabetes 15, 355–362. 10.1111/pedi.12092.

73. Carpenter, A.E., Jones, T.R., Lamprecht, M.R., Clarke, C., Kang, I.H., Friman, O., Guertin, D.A., Chang, J.H., Lindquist, R.A., Moffat, J., et al. (2006). CellProfiler: image analysis software for identifying and quantifying cell phenotypes. Genome Biology 7, R100. 10.1186/gb-2006-7-10-r100.

74. Looney, B.M., Xia, C.-Q., Concannon, P., Ostrov, D.A., and Clare-Salzler, M.J. (2015). Effects of Type 1 Diabetes-Associated IFIH1 Polymorphisms on MDA5 Function and Expression. Curr Diab Rep 15, 96. 10.1007/s11892-015-0656-8.

75. Schindelin, J., Arganda-Carreras, I., Frise, E., Kaynig, V., Longair, M., Pietzsch, T., Preibisch, S., Rueden, C., Saalfeld, S., Schmid, B., et al. (2012). Fiji: an open-source platform for biological-image analysis. Nat Methods 9, 676–682. 10.1038/nmeth.2019.

76. Berg, S., Kutra, D., Kroeger, T., Straehle, C.N., Kausler, B.X., Haubold, C., Schiegg, M., Ales, J., Beier, T., Rudy, M., et al. (2019). ilastik: interactive machine learning for (bio)image analysis. Nat Methods 16, 1226–1232. 10.1038/s41592-019-0582-9.

77. Windhager, J., Zanotelli, V.R.T., Schulz, D., Meyer, L., Daniel, M., Bodenmiller, B., and Eling, N. (2023). An end-to-end workflow for multiplexed image processing and analysis. Nat Protoc 18, 3565–3613. 10.1038/s41596-023-00881-0.

78. Virshup, I., Bredikhin, D., Heumos, L., Palla, G., Sturm, G., Gayoso, A., Kats, I., Koutrouli, M., Berger, B., Pe’er, D., et al. (2023). The scverse project provides a computational ecosystem for single-cell omics data analysis. Nat Biotechnol 41, 604–606. 10.1038/s41587-023-01733-8.

79. Paszke, A., Gross, S., Massa, F., Lerer, A., Bradbury, J., Chanan, G., Killeen, T., Lin, Z., Gimelshein, N., Antiga, L., et al. (2019). PyTorch: An Imperative Style, High-Performance Deep Learning Library. arXiv.org. https://arxiv.org/abs/1912.01703v1.

80. Chevrier, S., Crowell, H.L., Zanotelli, V.R.T., Engler, S., Robinson, M.D., and Bodenmiller, B. (2018). Compensation of Signal Spillover in Suspension and Imaging Mass Cytometry. Cell Syst 6, 612–620.e5. 10.1016/j.cels.2018.02.010.

81. Levine, J.H., Simonds, E.F., Bendall, S.C., Davis, K.L., Amir, E.D., Tadmor, M.D., Litvin, O., Fienberg, H.G., Jager, A., Zunder, E.R., et al. (2015). Data-Driven Phenotypic Dissection of AML Reveals Progenitor-like Cells that Correlate with Prognosis. Cell 162, 184–197. 10.1016/j.cell.2015.05.047.

82. Traag, V.A., Waltman, L., and van Eck, N.J. (2019). From Louvain to Leiden: guaranteeing well-connected communities. Sci Rep 9, 5233. 10.1038/s41598-019-41695-z.

83. Eling, N., Damond, N., Hoch, T., and Bodenmiller, B. (2021). cytomapper: an R/Bioconductor package for visualization of highly multiplexed imaging data. Bioinformatics 36, 5706–5708. 10.1093/bioinformatics/btaa1061.

84. Meyer, L., Eling, N., and Bodenmiller, B. (2024). cytoviewer: an R/Bioconductor package for interactive visualization and exploration of highly multiplexed imaging data. BMC Bioinformatics 25, 9. 10.1186/s12859-023-05546-z.

85. Breiman, L. (2001). Random Forests. Machine Learning 45, 5–32. 10.1023/A:1010933404324.

86. Sofroniew, N., Lambert, T., Bokota, G., Nunez-Iglesias, J., Sobolewski, P., Sweet, A., Gaifas, L., Evans, K., Burt, A., Doncila Pop, D., et al. (2024). napari: a multi-dimensional image viewer for Python. Version v0.5.4 (Zenodo). 10.5281/zenodo.13863809.

87. Kuznetsova, A., Brockhoff, P.B., and Christensen, R.H.B. (2017). lmerTest Package: Tests in Linear Mixed Effects Models. Journal of Statistical Software 82, 1–26. 10.18637/jss.v082.i13.

88. Robinson, M.D., McCarthy, D.J., and Smyth, G.K. (2010). edgeR: a Bioconductor package for differential expression analysis of digital gene expression data. Bioinformatics 26, 139–140. 10.1093/bioinformatics/btp616.

89. Angerer, P., Haghverdi, L., Büttner, M., Theis, F.J., Marr, C., and Buettner, F. (2016). destiny: diffusion maps for large-scale single-cell data in R. Bioinformatics 32, 1241–1243. 10.1093/bioinformatics/btv715.

90. Van den Berge, K., Roux de Bézieux, H., Street, K., Saelens, W., Cannoodt, R., Saeys, Y., Dudoit, S., and Clement, L. (2020). Trajectory-based differential expression analysis for single-cell sequencing data. Nat Commun 11, 1201. 10.1038/s41467-020-14766-3.

91. Catena, R., Özcan, A., Jacobs, A., Chevrier, S., and Bodenmiller, B. (2016). AirLab: a cloud-based platform to manage and share antibody-based single-cell research. Genome Biology 17, 142. 10.1186/s13059-016-1006-0.

92. Wolf, F.A., Angerer, P., and Theis, F.J. (2018). SCANPY: large-scale single-cell gene expression data analysis. Genome Biology 19, 15. 10.1186/s13059-017-1382-0.

93. Sella, F., Raz, G., and Cohen Kadosh, R. (2021). When randomisation is not good enough: Matching groups in intervention studies. Psychon Bull Rev 28, 2085–2093. 10.3758/s13423-021-01970-5.

94. Greenwald, N.F., Miller, G., Moen, E., Kong, A., Kagel, A., Dougherty, T., Fullaway, C.C., McIntosh, B.J., Leow, K.X., Schwartz, M.S., et al. (2022). Whole-cell segmentation of tissue images with human-level performance using large-scale data annotation and deep learning. Nat Biotechnol 40, 555–565. 10.1038/s41587-021-01094-0.

95. Haghverdi, L., Lun, A.T.L., Morgan, M.D., and Marioni, J.C. (2018). Batch effects in single-cell RNA-sequencing data are corrected by matching mutual nearest neighbors. Nat Biotechnol 36, 421–427. 10.1038/nbt.4091.

96. Wright, M.N., and Ziegler, A. (2017). ranger: A Fast Implementation of Random Forests for High Dimensional Data in C++ and R. Journal of Statistical Software 77, 1–17. 10.18637/jss.v077.i01.

97. Kmieciak, M., Gowda, M., Graham, L., Godder, K., Bear, H.D., Marincola, F.M., and Manjili, M.H. (2009). Human T cells express CD25 and Foxp3 upon activation and exhibit effector/memory phenotypes without any regulatory/suppressor function. Journal of Translational Medicine 7, 89. 10.1186/1479-5876-7-89.

98. Controlling the False Discovery Rate: A Practical and Powerful Approach to Multiple Testing - Benjamini - 1995 - Journal of the Royal Statistical Society: Series B (Methodological) - Wiley Online Library https://rss.onlinelibrary.wiley.com/doi/10.1111/j.2517-6161.1995.tb02031.x.

