## Supplementary material for "Imaging mass cytometry reveals early β-cell dysfunction and changes in immune signatures during type 1 diabetes progression in human pancreata": Document S1 (Figure S1-5)

#### Supplementary Figures

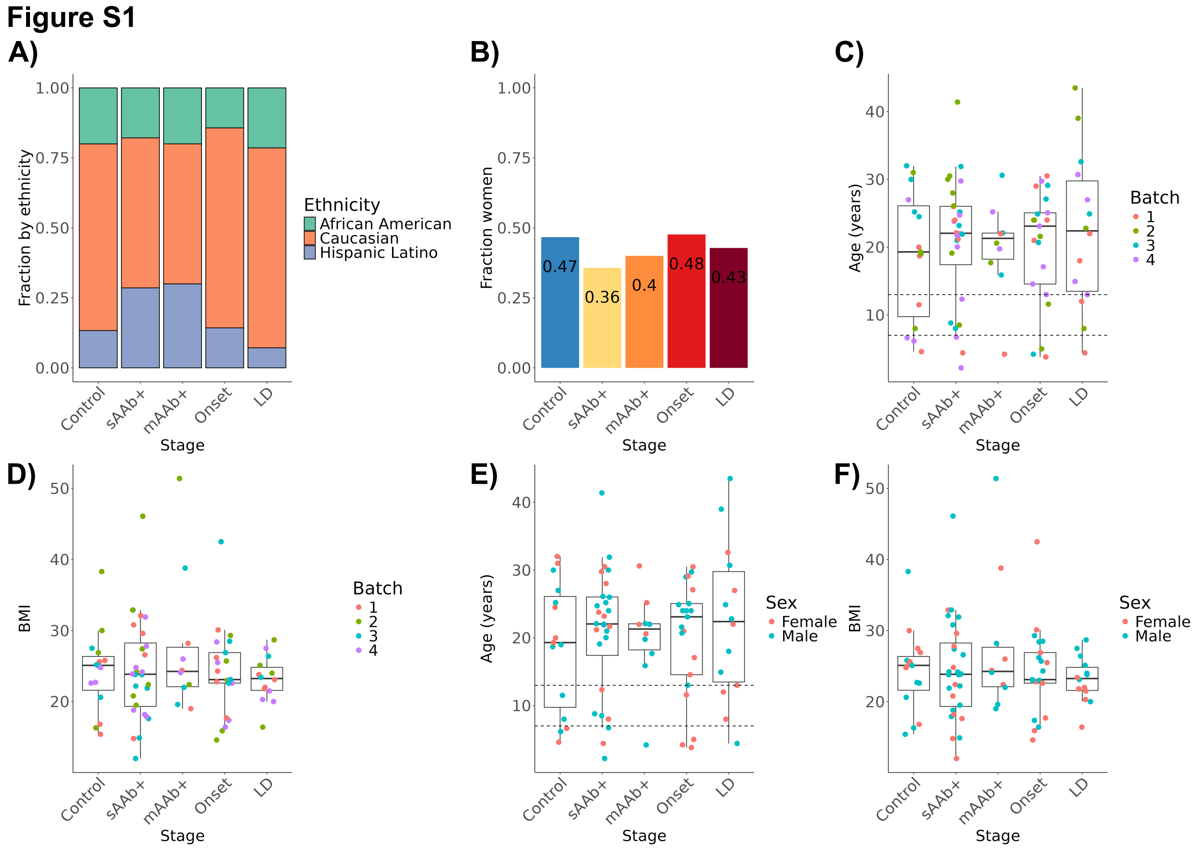


**Figure S1:** *IMC cohort is matched by BMI, age and gender.* **A:** Fraction of donors by ethnicity and disease stage. **B:** Fraction of donors by sex and disease stage. **C and D:** Data was acquired in four batches, and donors were matched by age **(C)** and BMI **(D)** across batches. **E and F:** Donors were matched by age, sex and BMI across disease stages.

#### Figure S2

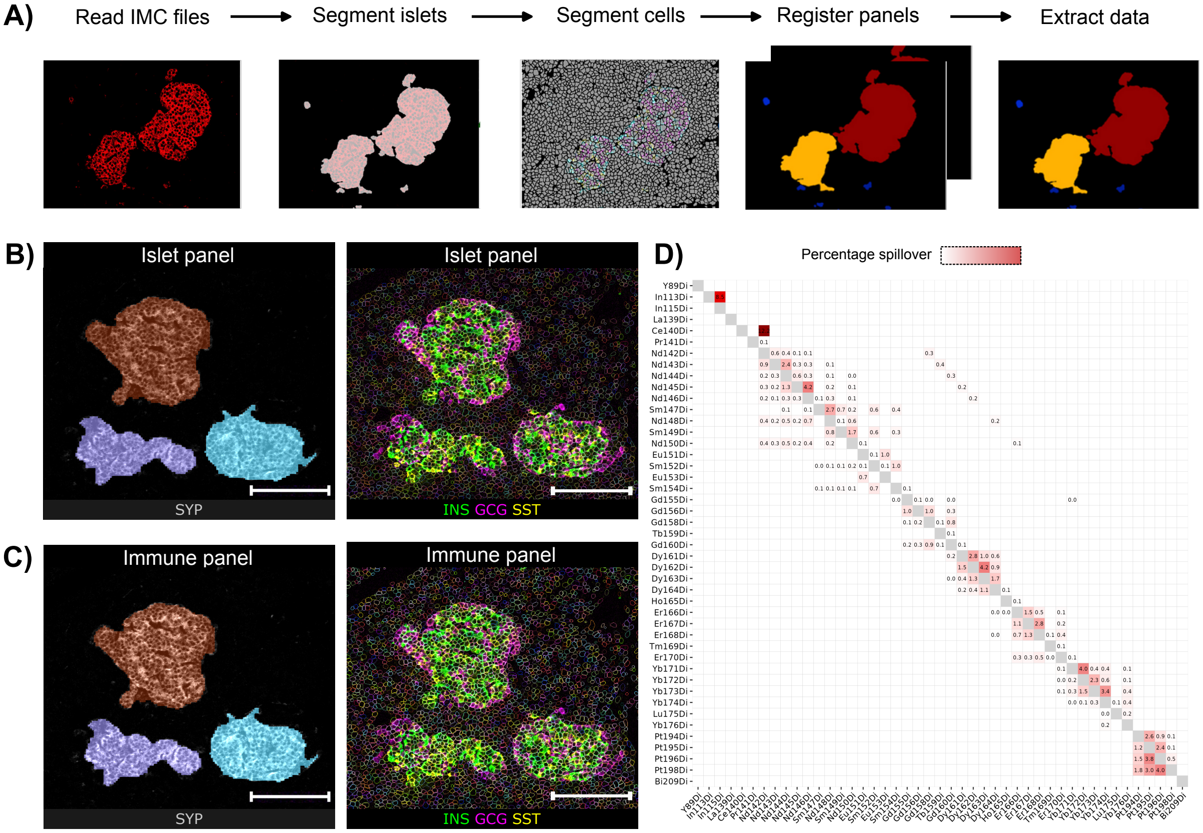


**Figure S2:** *IMC pre-processing workflow.* **A:** Schematic of the implemented pre-processing pipeline. **B)** Islet masks of acquired images from serial sections stained with the islet and immune panel. Scale bar is 100 μm. **C)** Cell masks of acquired images from serial sections stained with the islet and immune panel. Scale bar is 100 μm. **D)** Representative spillover matrix.

#### Figure S3

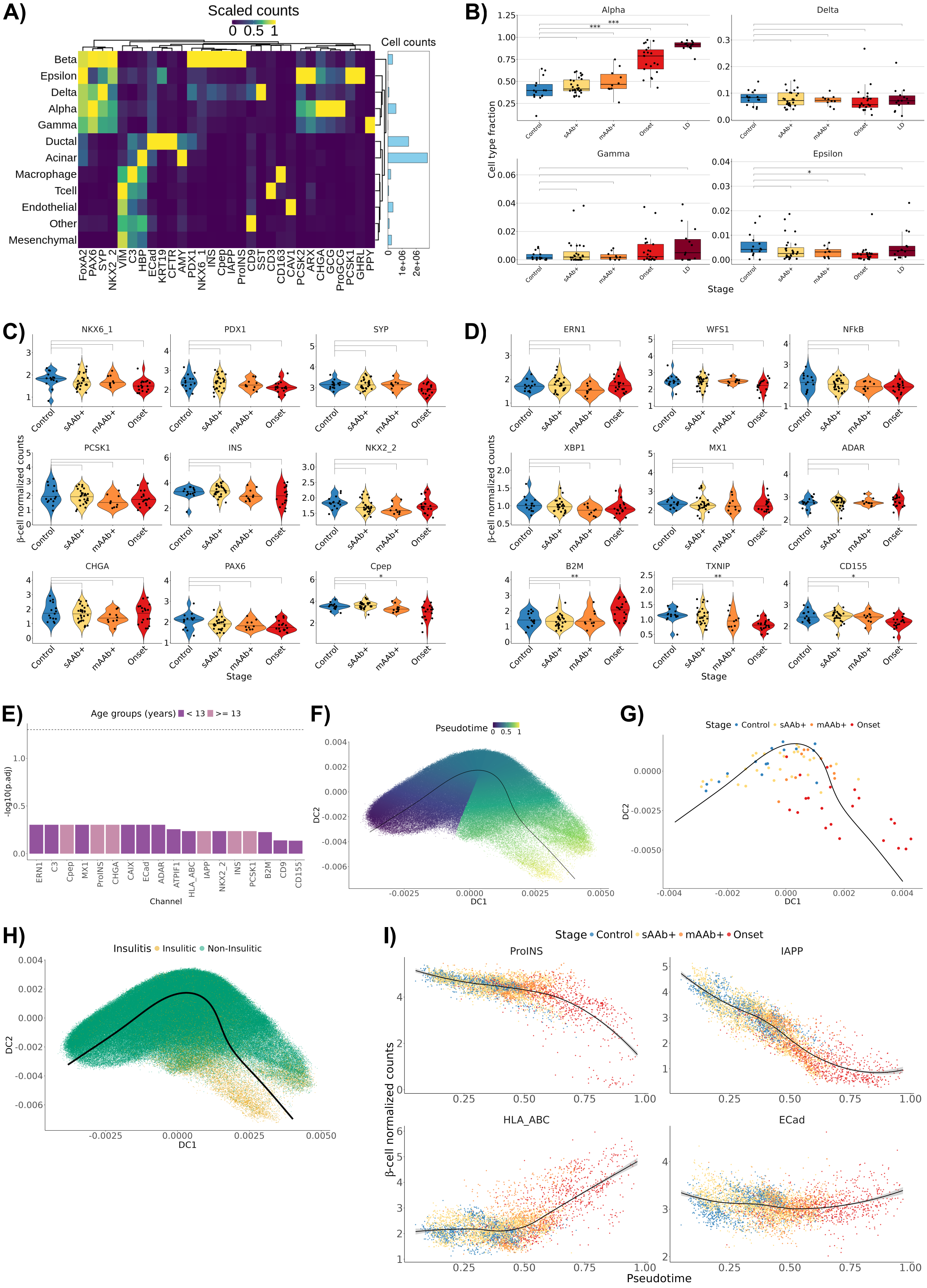

A)
Scaled counts
0 0.5 1
Cell counts
Beta
Epsilon
Delta
Alpha
Gamma
Ductal
Acinar
Macrophage
Tcell
Endothelial
Other
Mesenchymal
B)
Alpha
Delta
Gamma
Epsilon
Cell type fraction
Stage
C)
D)
β-cell normalized counts
Stage
E)
Age groups (years) < 13 >= 13
-log10(p.adj)
Channel
F)
Pseudotime
DC1
DC2
G)
Stage Control sAAb+ mAAb+ Onset
H)
Insulitis Insulitic Non-Insulitic
I)
ProINS
IAPP
HLA_ABC
ECad
Pseudotime

**Figure S3:** *β-cell expression profiles change with disease stage*. **A:** Heatmap of lineage and functional marker expression in annotated cell types. The barplot indicates the total counts of the given cell type. **B:** Fractions of α-, δ-, ε-, and γ-cells per islet in control and disease stage groups. Each dot indicates data from one donor averaged across ROIs. **C and D:** The asinh-normalized counts of **(C)** β-cell lineage and islet markers and **(D)** functional markers of the IFN-response (MX1, ADAR), ER stress (ERN1, WFS1, XBP1), inflammation signaling (NF-kB, CD155), redox signaling (TXNIP), and B2M subunit of the MHC-I complex across progression. Each dot indicates data from one donor with expression averaged across ROIs. **E:** Significance of expression of indicated makers in a linear mixed-effects model comparing samples from donors <13 years and ≥13 years for Onset T1D donors. Coloring indicates higher expression levels in the <13 years group (e.g., MX1) or ≥13 years group (e.g., Cpep). Only channels with an unadjusted p-value <0.5 are shown; no tests were significant. Y-axis is the -log10 adjusted p-values. **F:** Diffusion map-embedded β-cells colored by their respective pseudotime as predicted by the *slingshot* trajectory. **G:** Inferred pseudotime averaged by donor and visualized on the diffusion map shown in panel F. **H:** Diffusion-map embedded β-cells colored if they are from an insulitic (gold) or non-insulitic (green) islets. **I**: Expression of the indicated markers averaged by donor and visualized along the inferred pseudotime. Tests: Linear mixed-effects models with random intercept by organ donor were used to compare differential expression between disease stages against control donors. Differential abundance between stages was computed using *edgeR* and comparison against control donors. * *P* <0.05; ** *P* <0.01; *** *P* <0.001 for all comparisons.

### Figure S4

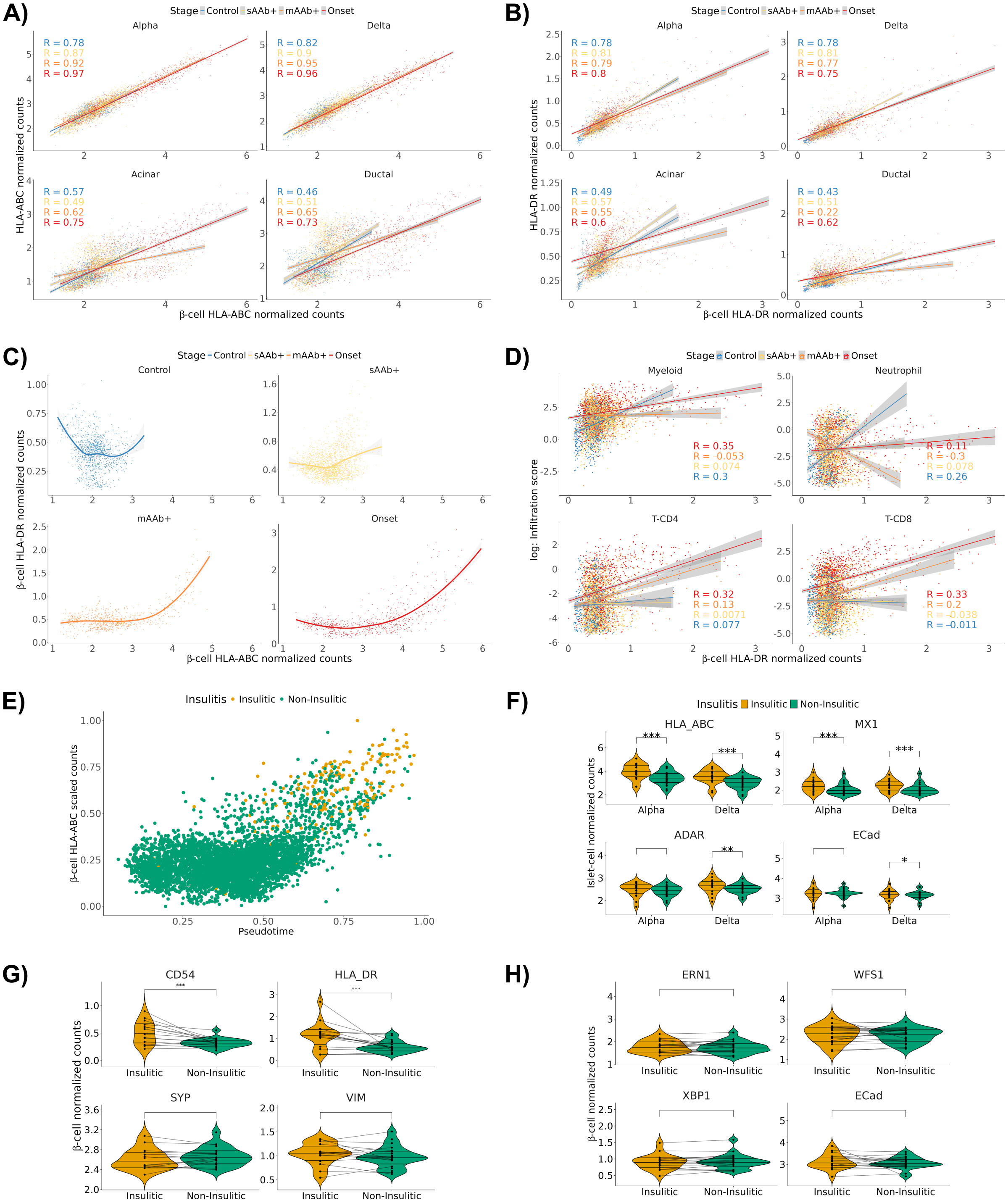

A)
Stage Control sAAb+ mAAb+ Onset
Alpha
Delta
Acinar
Ductal
HLA-ABC normalized counts
β-cell HLA-ABC normalized counts
B)
Stage Control sAAb+ mAAb+ Onset
Alpha
Delta
Acinar
Ductal
HLA-DR normalized counts
β-cell HLA-DR normalized counts
C)
Stage Control sAAb+ mAAb+ Onset
Control
sAAb+
mAAb+
Onset
β-cell HLA-DR normalized counts
β-cell HLA-ABC normalized counts
D)
Stage Control sAAb+ mAAb+ Onset
Myeloid
Neutrophil
T-CD4
T-CD8
log: Infiltration score
β-cell HLA-DR normalized counts
E)
Insulitis Insulitic Non-Insulitic
β-cell HLA-ABC scaled counts
Pseudotime
F)
Insulitis Insulitic Non-Insulitic
HLA_ABC
MX1
ADAR
ECad
Islet-cell normalized counts
Alpha
Delta
G)
CD54
HLA_DR
SYP
VIM
β-cell normalized counts
Insulitic
Non-Insulitic
H)
ERN1
WFS1
XBP1
ECad
β-cell normalized counts
Insulitic
Non-Insulitic

**Figure S4:** *HLA-DR is upregulated in insulitic islets.* **A and B:** Correlation of (A) HLA-ABC or (B) HLA-DR expression in the indicated cell types with expression in β-cells. Each dot is the average expression per ROI. Linear regression lines were fitted and colored per disease stage. **C:** Correlation between HLA-DR and HLA-ABC expression in β-cells in control and diseased tissues. Each dot is the average expression per ROI. LOWESS curves were fitted by disease stage. **D:** Correlations between cell density and HLA-DR expression in β-cells for the indicated immune cell types and disease stages. Each dot represents the averages per ROI. Linear regression lines were fitted by disease stage. **E:** HLA-ABC expression in β-cells as a function of pseudotime. Each dot is the average expression per ROI; values were min-max scaled. Gold-colored dots indicate ROIs containing insulitic islets. **F:** Violin plots of mean expression levels of the indicated markers in α- and δ-cells between ROIs containing insulitic and non-insulitic islets. The same ROIs were analyzed as in Figure 2F. Each dot indicates the average expression per sample across insulitic or non-insulitic ROIs. The minor change in E-Cad demonstrates that changes are not caused by general trends in protein levels. Linear mixed effects models were fit to determine significance. **G and H**: Violin plots of mean expression levels of the indicated markers in β-cells between ROIs containing insulitic and non-insulitic islets. Markers were stained with **(G)** the islet panel or **(H)** the immune panel. Lines connect insulitic and non-insulitic ROIs from the same donor. The minor expression changes in VIM and E-Cad demonstrate that changes are not caused by faulty β-immune cell segmentation or general trends in protein levels. Tests: * $P < 0.05$; ** $P < 0.01$; *** $P < 0.001$ for all comparisons**.** Linear mixed effects models were fit to determine significance between insulitic and non-insulitic ROIs.

#### Figure S5

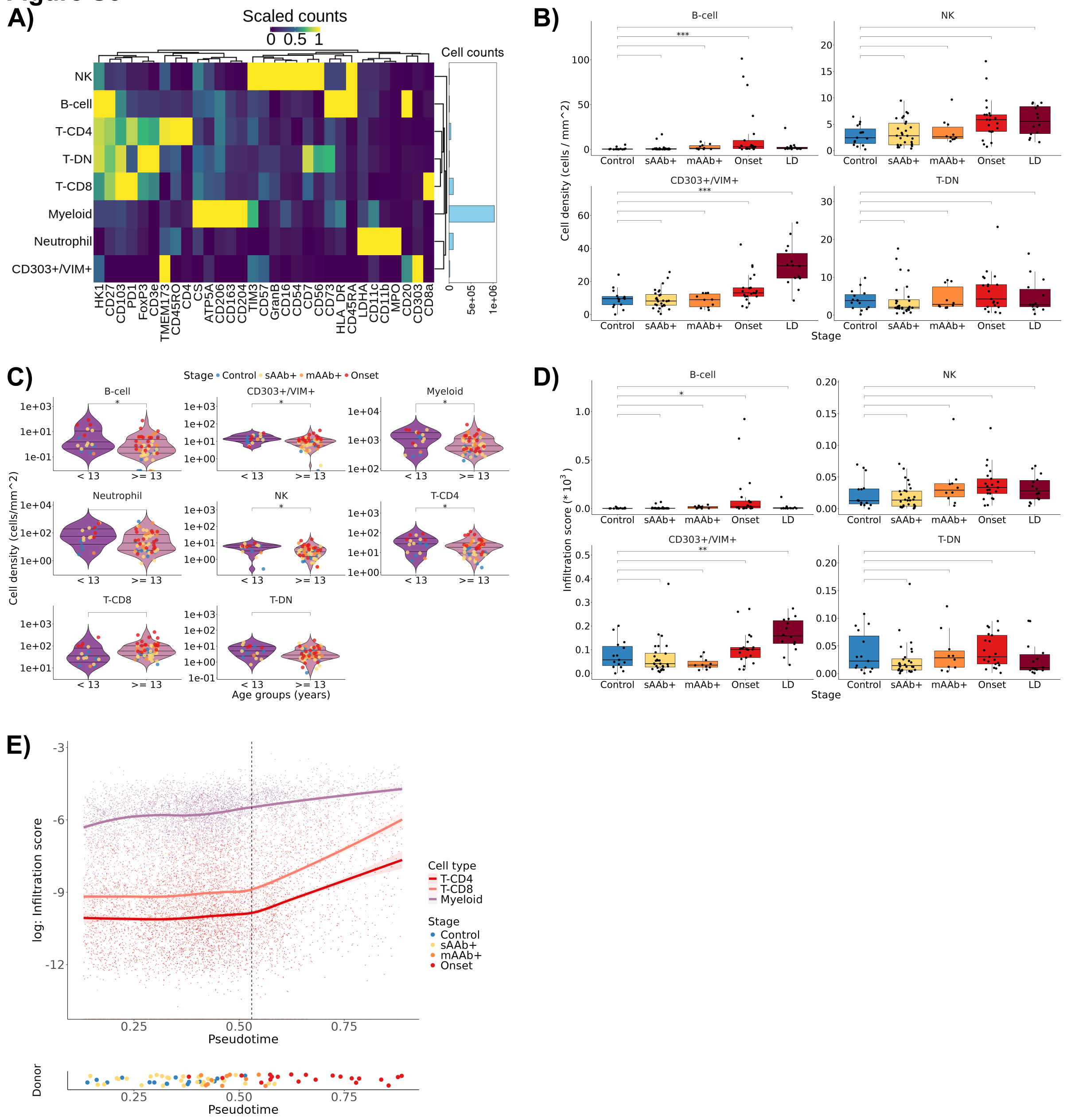


***Figure S5:*** *Immune cells infiltrate islets during disease progression.* **A:** Lineage marker expression by all annotated immune cell types. The barplot indicates total cell counts. **B:** Mean densities of the indicated cell types in control and disease tissue samples. Each dot indicates the mean cell density per sample averaged across ROIs. **C:** Violin plots of mean densities of the indicated cell types by donor age. Each dot is colored by disease stage and indicates the cell density per sample averaged across ROIs. Significance of difference between age groups was calculated by a linear mixed-effects model adjusted for disease stage. **D:** Mean infiltration scores for control and each disease stage. Each dot indicates data from one donor with infiltration score averaged across ROIs. **E:** Immune cell infiltration scores for the indicated cell types along pseudotime. Dots and regression lines are colored by cell type. Tests: * $P$ <0.05; ** $P$ <0.01; *** $P$ <0.001 for all comparisons. Significance was determined for each stage by testing for differential abundance against controls using *edgeR* and fitting linear mixed effects model between disease stages and controls.
