## Supplementary material for "Imaging mass cytometry reveals early β-cell dysfunction and changes in immune signatures during type 1 diabetes progression in human pancreata": Document S2 (Figure S6-10)

### Supplementary Figures S6-10

### Figure S6

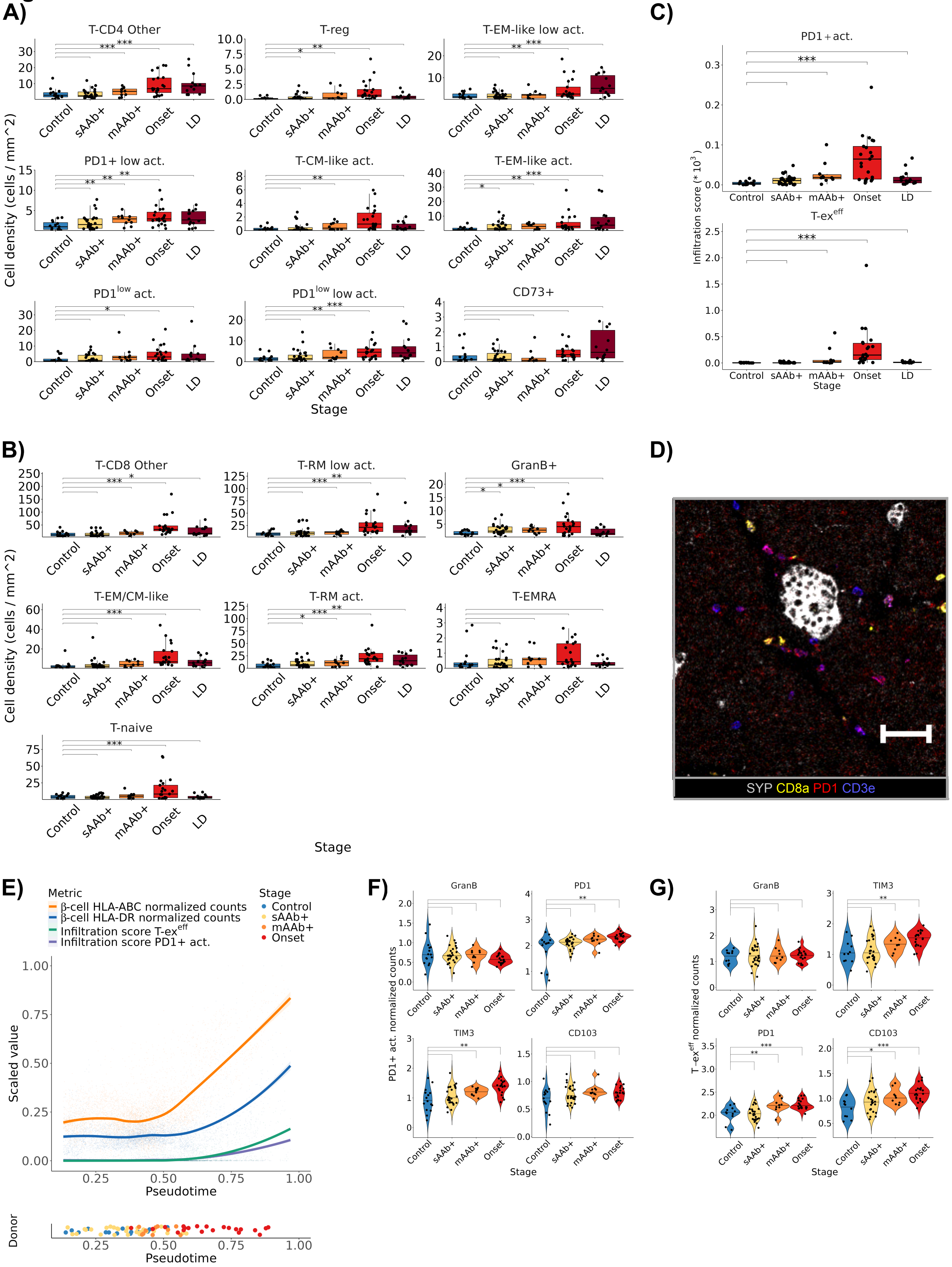

A)
T-CD4 Other
T-reg
T-EM-like low act.
PD1+ low act.
T-CM-like act.
T-EM-like act.
PD1low act.
PD1low low act.
CD73+
Cell density (cells / mm^2)
Control
sAAb+
mAAb+
Onset
LD
Stage
B)
T-CD8 Other
T-RM low act.
GranB+
T-EM/CM-like
T-RM act.
T-EMRA
T-naive
Cell density (cells / mm^2)
Stage
C)
PD1+act.
T-exeff
Infiltration score (* 10^3)
Stage
D)
SYP CD8a PD1 CD3e
E)
Metric
β-cell HLA-ABC normalized counts
β-cell HLA-DR normalized counts
Infiltration score T-exeff
Infiltration score PD1+ act.
Stage
Control
sAAb+
mAAb+
Onset
Scaled value
Pseudotime
Donor
F)
GranB
PD1
TIM3
CD103
PD1+ act. normalized counts
Stage
G)
GranB
TIM3
PD1
CD103
T-exeff normalized counts
Stage

***Figure S6:*** *Cell density and infiltration scores of T-CD4 and T-CD8 cell subtypes.* **A:** Boxplots of mean densities of the indicated T-CD4 **(A)** and T-CD8 **(B)** cell subtypes by disease stage. Each dot indicates data from one donor with cell density averaged across ROIs. **C:** Boxplots of mean infiltration scores of PD1+ act. cells (upper panel) and T-$ex^{eff}$ cells (lower panel) by disease stage. Each dot indicates data from one donor with cell infiltration score averaged across ROIs. **D:** Image of a sAAb+ donor stained for islets (SYP; gray), T cells (CD3e; blue), CD8a (yellow), and PD1 (red). Image depicts presence of PD1+ act. cells ($CD3e^{+}PD1^{+}$; purple) in islet-proximal exocrine tissue, but not islets in sAAb+ donors. PD1+ signal intensity is lower in comparison to islet-infiltrating cells. **E:** Scaled infiltration scores of PD1+ act. cells and T-$ex^{eff}$ cells along pseudotime per ROI. Infiltration scores are aligned with β-cell HLA-ABC and HLA-DR min-maxed values. Regression lines were fitted with LOWESS, with the shaded areas indicating the confidence interval of 95%. Average pseudotime per donors is indicated in the lower panel and are colored by disease stage. **F and G:** Expression levels of the indicated markers in PD1+ act. cells **(F)** and T-$ex^{eff}$ cells **(G)** across disease stages. Each dot indicates data from one donor with marker expression averaged across ROIs containing β-cells. Tests: * $P$ <0.05; ** $P$ <0.01; *** $P$ <0.001 for all comparisons. Significance was determined for each stage by testing for differential abundance against controls using *edgeR* or by fitting a linear mixed-effects model to test for increased infiltration score or differential expression against controls.

### Figure S7

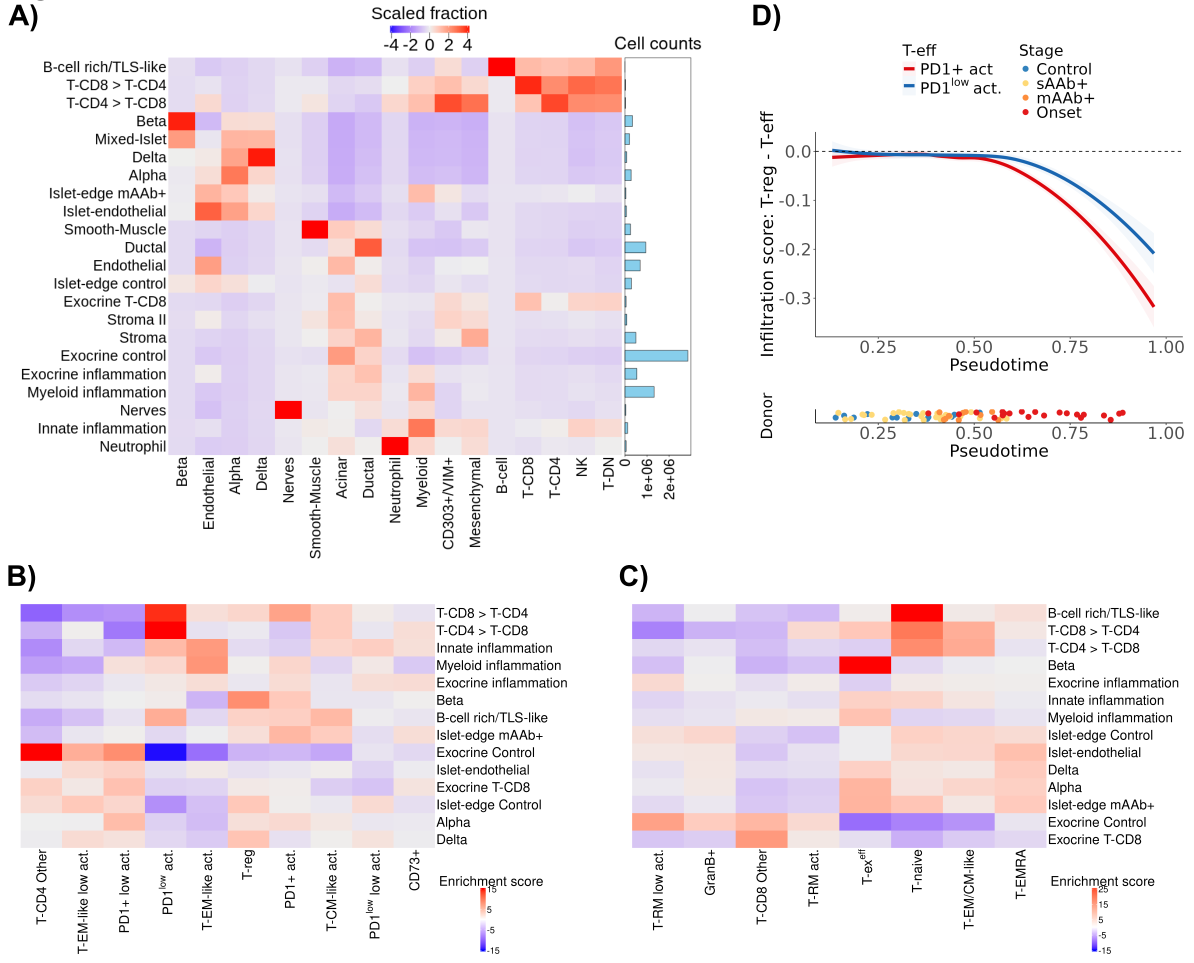


**Figure S7:** *Enrichment of exhausted-like T cells in peri-islet and β-cell cellular neighborhoods.* **A:** The cell compositions of aggregated cellular neighborhoods (CNs). Columns are scaled by cell type fractions in the indicated CNs. Red indicates higher fractions, blue lower fractions. Barplot to the right indicates the total numbers of cells per CN. **B and C:** Enrichment of the indicated T-CD4 subtypes **(B)** and T-CD8 subtypes **(C)** in informative CNs. Red indicates enrichment, blue indicates absence. Matrix elements is the enrichment score ($\chi^2$-test residuals). **D:** Infiltration scores of distinct T-CD4 effector cells (T-eff) (PD1+ act., PD1$^{low}$ act.) along pseudotime per ROI. Infiltration scores were subtracted against T-reg infiltration scores. Negative values thus indicate higher presence of T-eff than T-reg cells in islets. Regression lines were fitted with LOWESS, with the shaded areas indicating the confidence interval of 95%. Average pseudotime per donor is indicated by the x-axis position of the labeled dots in the lower panel.

### Figure S8

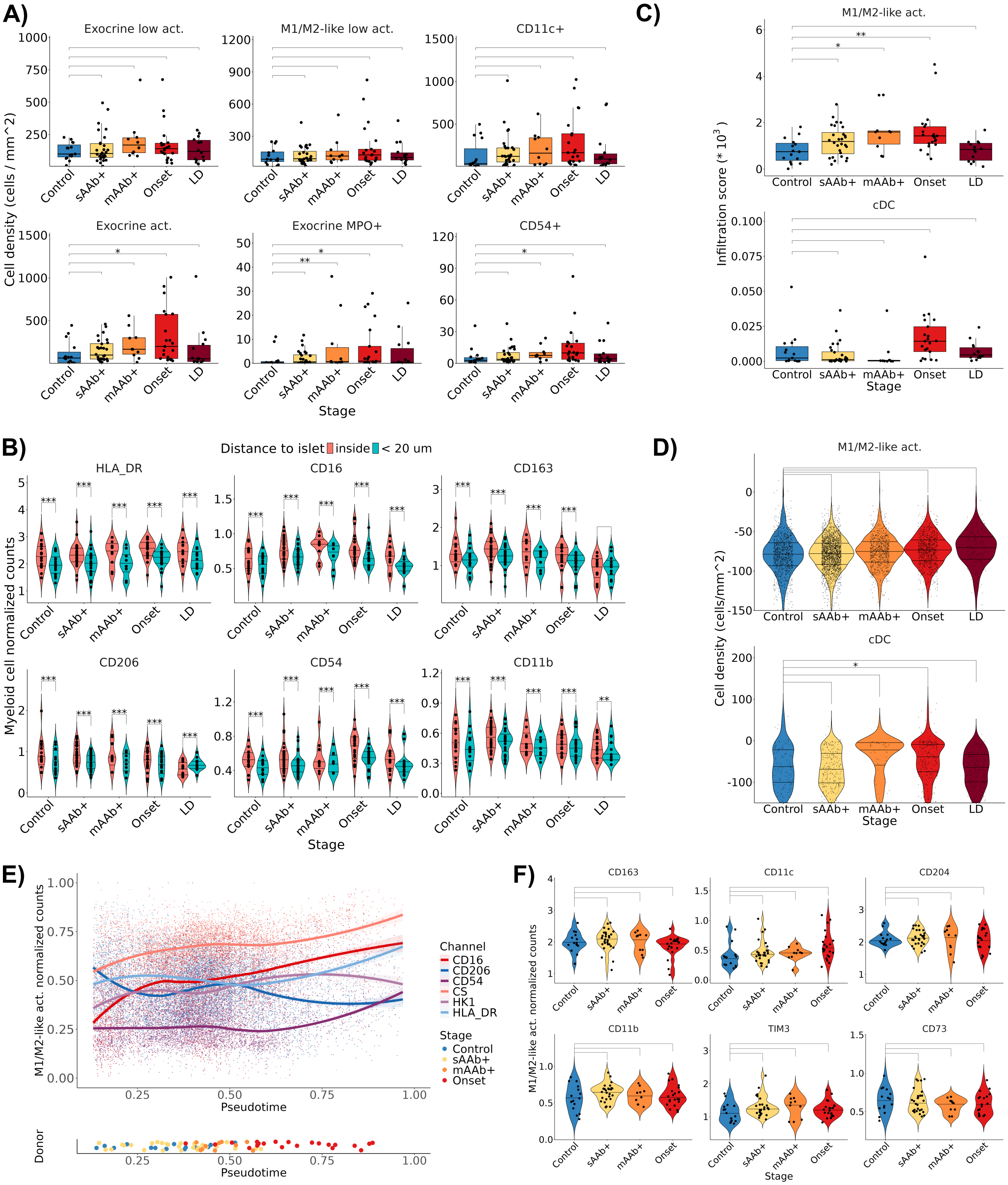

A)
Exocrine low act.
M1/M2-like low act.
CD11c+
Exocrine act.
Exocrine MPO+
CD54+
Cell density (cells / mm^2)
Control
sAAb+
mAAb+
Onset
LD
Stage
B)
Distance to islet
inside
< 20 um
HLA_DR
CD16
CD163
CD206
CD54
CD11b
Myeloid cell normalized counts
C)
M1/M2-like act.
cDC
Infiltration score (* 10^3)
D)
Cell density (cells/mm^2)
E)
M1/M2-like act. normalized counts
Pseudotime
Channel
CD16
CD206
CD54
CS
HK1
HLA_DR
Donor
F)
CD163
CD11c
CD204
CD11b
TIM3
CD73
M1/M2-like act. normalized counts

***Figure S8:*** *cDCs and activated M1/M2-like macrophages are attracted to islets in T1D pancreas.* **A and C:** Boxplots comparing **(A)** mean densities and **(C)** mean infiltration scores of the indicated myeloid cell subtypes between different disease stages. Each dot indicates the cell density or infiltration score for one donor averaged across ROIs. **B:** Expression of key functional and lineage markers of islet and peri-islet myeloid cells across disease stages. Each dot indicates data from one donor with expression averaged across ROIs. Significance was tested between islet and peri-islet myeloid expression by fitting a linear mixed-effects model. **D:** Violin plots of mean distances of M1/M2-like act. macrophages and cDCs to islets across disease stages. Given different abundances, each dot indicates single-cell distance measurements for cDCs, and ROI-average distance measurements for M1/M2-like act. macrophages. Statistics were computed on the sample level by linear mixed-effects models. **E:** Scaled expression levels of key myeloid markers along the inferred β-cell pseudotime trajectory. **F:** Expression levels of the indicated myeloid and metabolic markers in M1/M2-like act. macrophages across disease stages. Each dot indicates data from one donor with marker expression averaged across ROIs. Only ROIs containing β-cells were considered. Tests: * *P* <0.05; ** *P* <0.01; *** *P* <0.001 for all comparisons. Significance was determined for each stage by testing for differential abundance against controls using *edgeR* and by testing for differential expression against controls by fitting a linear mixed-effects model.

#### Figure S9

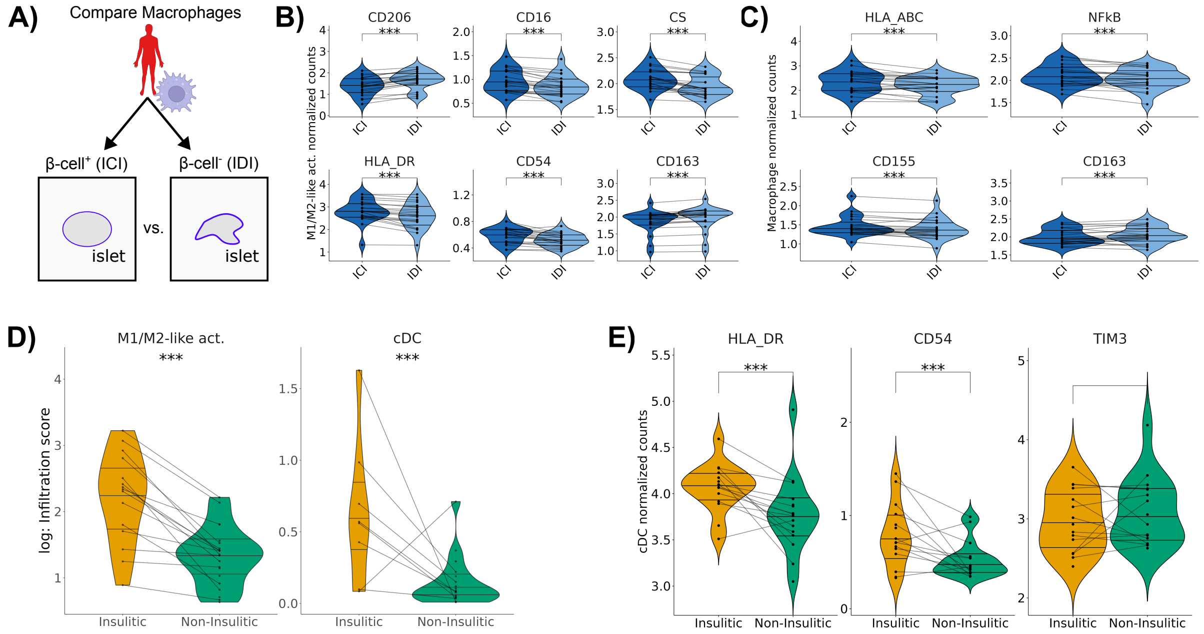


**Figure S9:** *Myeloid cells have pro-inflammatory signatures in insulitic and ICI ROIs.* **A:** Schematic of macrophage phenotypes comparison between ROIs that contain β-cells (ICIs) and those that do not (IDIs) from the same donor. **B and C:** Levels of key myeloid markers for **(B)** M1/M2-like act. macrophages and **(C)** peri-islet and islet macrophages (< 20 µm to the islet edge) between ICIs and IDIs of the same donor, as annotated by the islet panel. ICIs and IDIs from the same donor are linked by lines. Only Onset donors were considered. **D:** Violin plots of log-transformed infiltration scores of the indicated cell subtypes between insulitic (N=15 donors, N=97 ROIs) and non-insulitic ROIs (N=21 donors, i=813 ROIs). Insulitic and non-insulitic ROIs from the same donor are linked by lines. Only ICI ROIs of Onset T1D donors are considered. **E:** Violin plots of mean expression levels for the indicated markers in cDCs from insulitic (N=15 donors, i=97 ROIs) and non-insulitic ROIs (N=21 donors, i=813 ROIs). Insulitic and non-insulitic ROIs from the same donor are linked by lines. Only ICI ROIs of Onset donors are considered. Tests: * *P* <0.05; ** *P* <0.01; *** *P* <0.001 for all comparisons. Significance was determined by comparing donor-averaged myeloid subtype expression levels between ICIs and IDIs or non-insulitic and insulitic ROIs using a linear mixed-effects model.

#### Figure S10

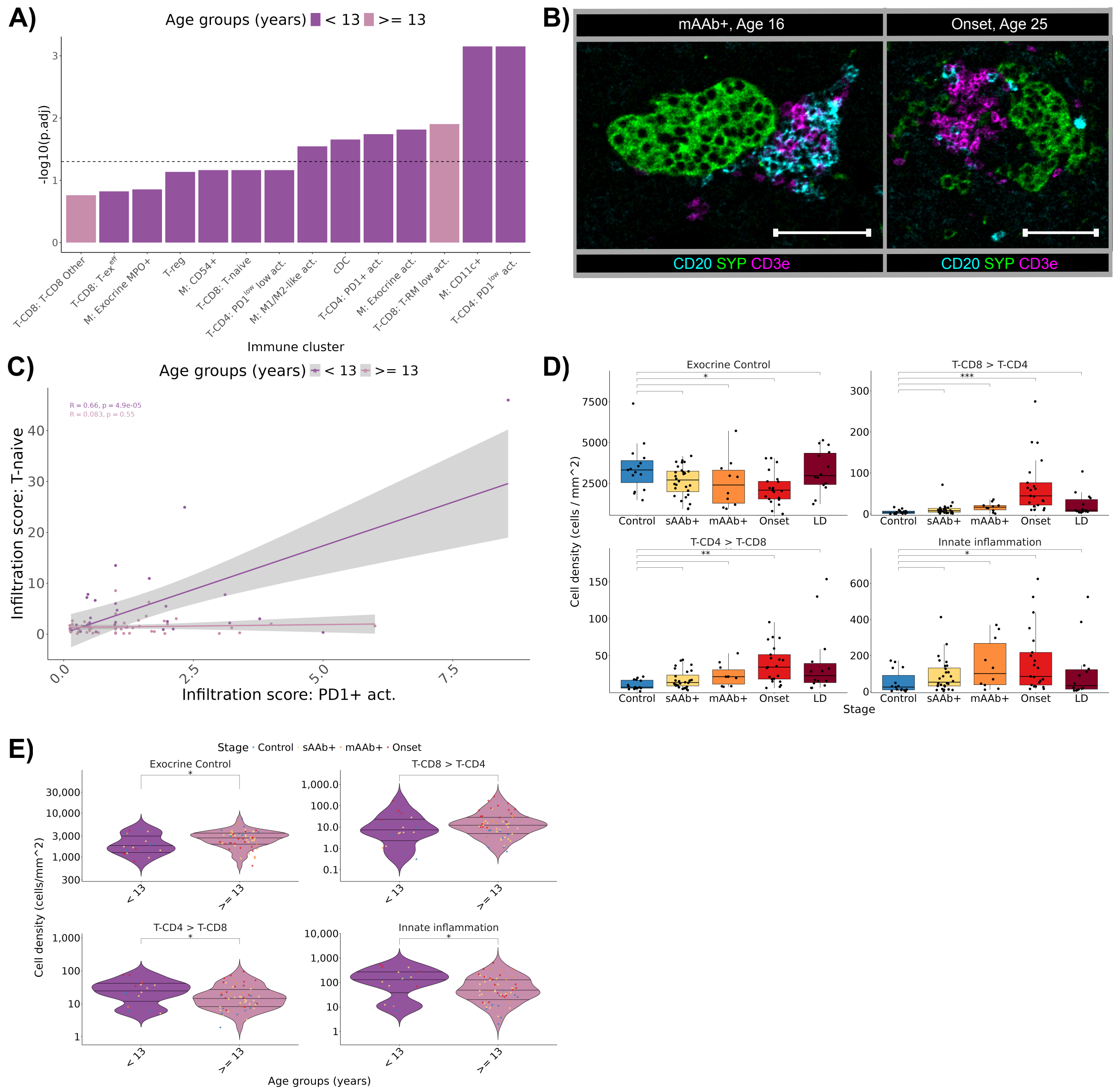


***Figure S10:*** *Spatial, progression, and age-associated immune cell motifs.* **A:** Regression analysis using a linear mixed-effects model of association of age with abundance of indicated immune cell subtypes. Density per immune cell subtype was calculated per ROI. The horizontal line indicates an adjusted p-value of 0.05. **B:** Micrographs of tissue from one mAAb+ (left) and one Onset donors (right) with significant $CD20^{+}$ B-cell (cyan) infiltration. **C:** Infiltration scores of naïve T-cells and PD1+ act. cells in insulitic islets. **D:** Boxplots of mean densities of the indicated CNs in control and diseased tissue. Each dot indicates data from one donor with cell density averaged across ROIs. **E:** Violin plots of mean density between younger (< 13 years) and older (≥13 years) donors for the indicated CNs. Each dot is colored by disease stage and denotes the mean cell density per sample across ROIs. Significance was calculated by a linear mixed-effects model, while adjusting for disease stage. Tests: * $P$ <0.05; ** $P$ <0.01; *** $P$ <0.001 for all comparisons.
